# Resting-state directed brain connectivity patterns in adolescents from source-reconstructed EEG signals based on information flow rate

**DOI:** 10.1101/608299

**Authors:** Dionissios T. Hristopulos, Arif Babul, Shazia Babul, Leyla R. Brucar, Naznin Virji-Babul

## Abstract

Quantifying the brain’s effective connectivity offers a unique window onto the causal architecture coupling the different regions of the brain. Here, we advocate a new, data-driven measure of directed (or effective) brain connectivity based on the recently developed information flow rate coefficient. The concept of the information flow rate is founded in the theory of stochastic dynamical systems and its derivation is based on first principles; unlike various commonly used linear and nonlinear correlations and empirical directional coefficients, the information flow rate can measure causal relations between time series with minimal assumptions. We apply the information flow rate to electroencephalography (EEG) signals in adolescent males to map out the directed, causal, spatial interactions between brain regions during resting-state conditions. To our knowledge, this is the first study of effective connectivity in the adolescent brain. Our analysis reveals that adolescents show a pattern of information flow that is strongly left lateralized, and consists of short and medium ranged bidirectional interactions across the frontal-central-temporal regions. These results suggest an intermediate state of brain maturation in adolescence.

## Introduction

The brain is a complex entity comprising widely distributed but highly interconnected regions, the dynamic interplay of which is essential for brain function. Establishing how activity is coordinated across these regions to give rise to organized (higher order) brain functions ranks as one of the key challenges in neuroscience. Various measures of brain connectivity are in use for this purpose as discussed in (*Friston, 1994*; *Horwitz, 2003*; *Sporns, 2011*; *Rubinov and Sporns, 2010*; *Friston, 2011*; *Cohen, 2014*) and references therein. Structural measures are based on confirmed anatomical connections between brain regions. Functional measures involve dynamically changing, linear or nonlinear, non-directional coefficients of statistical dependence (e.g., correlation, covariance, phase-locking values, coherence) that may appear between structurally unconnected regions. Effective brain connectivity measures capture directionally dependent interactions between different brain regions and aim to identify causal mechanisms in neural processing. In the following, we use the terms “effective” and “’directed” connectivity interchangeably. We refer readers to *Sakkalis (2011)* and *Bastos and Schoffelen (2015)* for recent reviews of functional and effective connectivity measures in the brain. Herein, we investigate effective connectivity patterns as revealed by electroencephalography (EEG) recordings (*Van de Ville et al., 2010*) of scalp electromagnetic fields following source-space reconstruction.

The multichannel EEG signals, which are thought to reflect activity in the underlying brain regions, offer a convenient window into the temporal dynamics of the corresponding brain-scale neuronal networks. EEG studies have been extensively used to infer the nature of the functional connectivity —-i.e. the linear or nonlinear statistical interdependence between the electrical activity in the different brain regions (*Stam and Van Straaten, 2012*) — during resting state or during task related activities. In this paper, we focus our attention on the former.

The resting-state or persistent background activity, previously dismissed as background noise, has been shown to comprise coherent patterns of functional connectivity and appears to play a critical role in mediating complex functions such as memory, language, speech and emotional states (*Raichle et al., 2001*; *Raichle and Mintun, 2006*). There has been considerable progress in mapping out the key resting-state functional brain networks as well as tracking how they change over development. These functional connectivity studies indicate that the resting-state brain networks are sparsely connected in childhood (*Fair et al., 2008*) and evolve towards increased connectivity in adolescence (*Smit et al., 2012*). However, a more complete description remains elusive. For one, very little is known about how information flows within these networks, and how these flow patterns change with maturation.

Several different approaches are in use for quantifying the brain’s effective connectivity. Structural approaches such as Structural Equation Modeling (SEM) (*McLntosh and Gonzalez-Lima, 1994*) and Dynamic Causal Modeling (*Friston et al., 2003*) involve a neuranatomical model of the brain and a connectivity model. Other measures are data-driven and involve a statistical model, such as Granger-causality-based methods (*Kamiński et al., 2001*; *Hesse et al., 2003*; *Roebroeck et al., 2005*; *Ding et al., 2006*; *Bressler and Seth, 2011*; *Seth et al., 2015*). A different data-driven approach involves information theoretic measures, like transfer entropy (*Schreiber, 2000*; *Vicente et al., 2011*) and partial directed coherence (*Baccalá and Sameshima, 2001*). Each approach has its advantages and disadvantages [see (*Lindquist, 2008*; *Liu and Aviyente, 2012*) and the Discussion section below] in terms of the assumptions involved and the computational effort required. The fact that all methods currently used make assumptions the validity of which has not been fully tested, leaves room for introducing new measures of effective connectivity (*Lindquist, 2008*). This motivates the investigation of new measures of directed connectivity.

We have two goals in this paper. Our first goal is to advocate a new measure of data-driven effective brain connectivity by applying the novel concept of information flow rate to EEG signals. This goal is motivated by the need to define measures of connectivity that are based on fewer or more suitable model assumptions than commonly used methods (*Lindquist, 2008*). The information flow rate has several desirable properties (as summarized below and elaborated in the Discussion section) which give it unique advantages for connectivity analysis compared with standard methods. To the best of our knowledge, our study is the first to apply the information flow rate to neuroscience data. Our second goal is to analyze EEG resting-state data from a group of healthy adolescents using the information flow rate, in order to identify connectivity patterns in the adolescent brain. Only one prior study that focuses on this age group is available in the literature, and the connectivity analysis in that study is carried out in sensor space (*Marshall et al., 2014*).

The information flow rate was developed by Liang using the concept of information entropy and the theory of dynamical systems (*Liang, 2008*, 2013b, *2014*, *2015*) and based on earlier work with Kleeman (*Liang and Kleeman, 2005*). While the the initial formulation of the information flow rate was derived for two-dimensional (bivariate) systems, *Liang (2016*, *2018*) recently showed that the formulation is also valid for *N*-dimensional systems as well. The Liang-Kleeman coefficient can measure the transfer of information between time series at different locations and thus between different brain regions. Unlike empirical measures of causality, e.g., transfer entropy and Granger causality, the information flow rate is derived from general, first-principles equations for the time evolution of stochastic dynamical systems (*Liang, 2016*, *2018*). Owing to its definition, which involves only the time series and their temporal derivatives (or their finite-difference approximations for discretely sampled systems), the information flow rate has computational advantages over other entropy-based measures such as transfer entropy, that require the estimation of additional information (e.g., conditional probabilities) from the data. In addition, the information flow rate concept does not require stationarity (*Liang, 2015*) or a specific model structure, and can also be applied to deterministic nonlinear systems (*Liang, 2016*). These are important advantages, since the EEG signals exhibit non-stationary features evidenced in transitions between quasi-stationary periods and nonlinear dynamic behavior (*Blanco et al., 1995*; *Kaplan et al., 2005*; *Klonowski, 2009*).

## Results

We set out to investigate patterns of resting-state effective connectivity in the brain of adolescent males, using source-reconstructed EEG signals (see Materials and methods). Our analysis of connectivity is based on the Liang-Kleeman information flow rate described in Box 1. The information flow rate measures the effect of a time series *i*, called *transmitter*, on a different time series *j*, called *receiver*. The indices *i* and *j* correspond to different brain source locations. In particular, we use a normalized version of the information flow rate which is better suited for ranking pair-wise information flow rates for an ensemble of *N*_*s*_ time series based on their relative impact on the *receiver time series* (see Materials and methods for details). Herein *N*_*s*_ = 15 corresponds to the numbers of source locations obtained by source-space reconstruction. The brief comment in Box 2 provides an intuitive understanding of causal relations in terms of the information flow rate.

### Box 1. The Liang-Kleeman information flow rate

Let 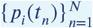 denote a collection of *N*_*s*_ time series at different brain source locations indexed by *i* = 1, …, *N_s_*. Herein, the term “time series” implies EEG-derived series of current dipole moments.

The *Liang-Kleeman coefficient T*_*i*→*j*_ measures the rate of information flow from the time series *i* to the time series *j* (where *j* ≠ *i*). *T*_*i→j*_ can be expressed in terms of *sample statistics* as follows (*Liang, 2014*)

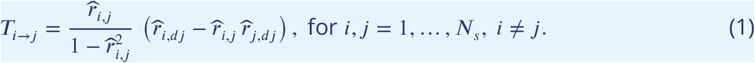

In the above, the *linear (Pearson) sample correlation coefficient* between the time series *p*_*i*_ and *p*_*j*_ is defined by

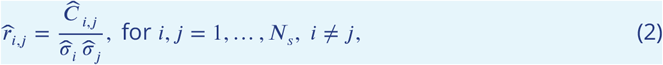

where 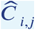 is the *sample cross-covariance* of the series *p*_*i*_ and *p*_*j*_, and 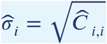 is the sample standard deviation of the series *p*_*i*_ (*i* = 1, …, *N_s_*). Both 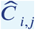 and 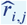(often used to measure functional connectivity) are non-directional and symmetric under the index interchange *i* ⇄ *j*. The sample cross-covariance 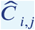 is defined by

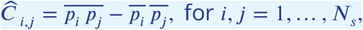

where the “overline” denotes the sample time average, i.e., 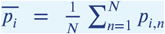 and 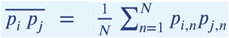. If *i* = *j* the above equation returns the variance of *p*_*i*_, i.e., 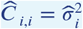.

The *cross-correlation coefficients* 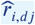, where *i, j* = 1, …, *N_s_*, in *Equation 1* involve the time series *p*_*i*_ and the *temporal derivative* 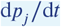 of the time series *p*_*j*_. These coefficients are expressed in terms of the respective covariances as follows

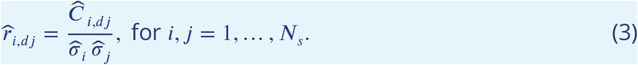

where 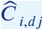 is the sample covariance of the time series *p*_*i*_ and the first derivative, 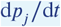, of the series *p*_*j*_. Due to the discrete nature of sampling, the first derivative 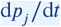 is unknown *a priori*. Hence, a finite difference approximation based on the Euler forward scheme, with a time step equal to *k*Δ*t*, is used, i.e.,

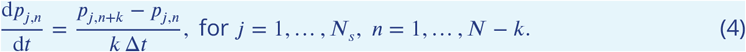

The differencing orders *k* = 1 and *k* = 2 are the two most common choices (Liang, 2013a) which we also consider herein.

Herein we refer to *p*_*i*_ as the *transmitter series* and to *p*_*j*_ as the *receiver series* with respect to *T*_*i*→*j*_. We adopt the term *transmitter* instead of “source” for the series that “sends” information in order to avoid confusion, since all the time series represent *current dipole moments* obtained from scalp EEG by means of *source reconstruction*.

### Box 2. Causality and the Liang-Kleeman coefficient

Consider two time series *p*_*i*_ and *p*_*j*_ where *i*, *j* = 1,…,*N*_*s*_ and *j* ≠ *i*. According to the Liang-Kleeman formalism which is based on the notion of information entropy, the series *p*_*j*_ has a causal effect on *p*_*i*_ if the rate of change of *p*_*i*_ depends on *p*_*j*_. Conversely, *p*_*i*_ has a causal effect on *p*_*j*_ if the rate of change of *p*_*j*_ depends on *p*_*i*_. Hence, the following four possibilities arise:

1. Neither *p*_*i*_ influences *p*_*j*_, nor *p*_*j*_ influences *p*_*i*_: *T*_*i*→*j*_ = *T*_*j*→*i*_ = 0:
2. Only *p*_*i*_ influences *p*_*j*_, but *p*_*j*_ does not influence *p*_*i*_: *T*_*i*→*j*_ ≠ 0, *T*_*j*→*i*_ = 0:
3. Only *p*_*j*_ influences *p*_*i*_, but *p*_*i*_ does not *p*_*i*_ influence *p*_*j*_: *T*_*i*→*j*_ = 0; *T*_*j*→*i*_ ≠ 0:
4. Both *p*_*i*_ and *p*_*j*_ influence each other: *T*_*i*→*j*_ ≠ 0, *T*_*j*→*i*_ ≠ 0.
5. *T*_*i*→*j*_ does not have a physical meaning and is thus undefined.

To quantify effective brain connectivity, we use the *normalized information flow rate r_i_*_→*j*_(defined by *Equation 9* in Material and methods). The time series that we analyze involve the magnitudes of fifteen *current dipole moments* per individual. These are obtained by means of source reconstruction of scalp EEG signals as described in Materials and Methods. We focus on the *normalized inter-dipole information flow rate* instead of the non-normalized *T*_*i*→*j*_, because we aim to capture interactions between brain regions that significantly affect the *receiver* region (denoted by the index *j*). The advantage of *τ*_*i*→*j*_ is its ability to measure the *relative importance* of causal relations (*Liang, 2015*).

We use the second-neighbor differencing scheme (i.e., *k* = 2, see Box 1) to calculate the information flow rates as suggested by *Liang (2014)*. We further comment on this choice in Materials and methods (section on the impact of differencing scheme).

Our analysis focuses on the *mean information flow rate* calculated over all the individuals in the study cohort, but we also explore variations of connectivity between individuals.

### Brain connectivity based on mean information flow rate

To study the information flow across brain regions we want to characterize connections that exhibit significant levels of activity (as measured by the information flow rate) over all the individuals. We do this using the *ensemble mean* of the normalized information flow rate *i* → *j* evaluated over the cohort of *L* = 32 individuals:

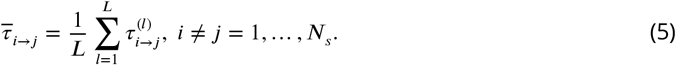

The top panel in *Figure 1* displays the patterns of the *mean information flow rate* 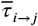. The variable 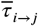 for all values of *transmitter i* and *receiver j* source locations is represented by an *N*_*s*_ × *N*_*s*_ matrix that represents all possible (i.e., 210) connections between sources. The number of possible connections is *N*_*s*_ × (*N*_*s*_ − 1) where *N*_*s*_ = 15 is the number of source dipoles. The value of a grid cell (L1, L2), determined by the label L1 on the vertical axis and the label L2 on the horizontal axis, represents information flow from dipole L1 to dipole L2. The matrix cells are colored according to the value of 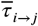: the values increase as the color changes from blue to red. The cells along the main diagonal are not colored, indicating that the information flow rate from *i* → *j* is only defined if *i* ≠ *j*. The color pattern (thus, also the matrix 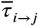) is *asymmetric* along the main diagonal. This asymmetry reflects the directionality of the information flow rate, i.e., the fact that *τ*_*i*→*j*_ is in general different than *τ*_*j*→*i*_.

**Figure 1.**
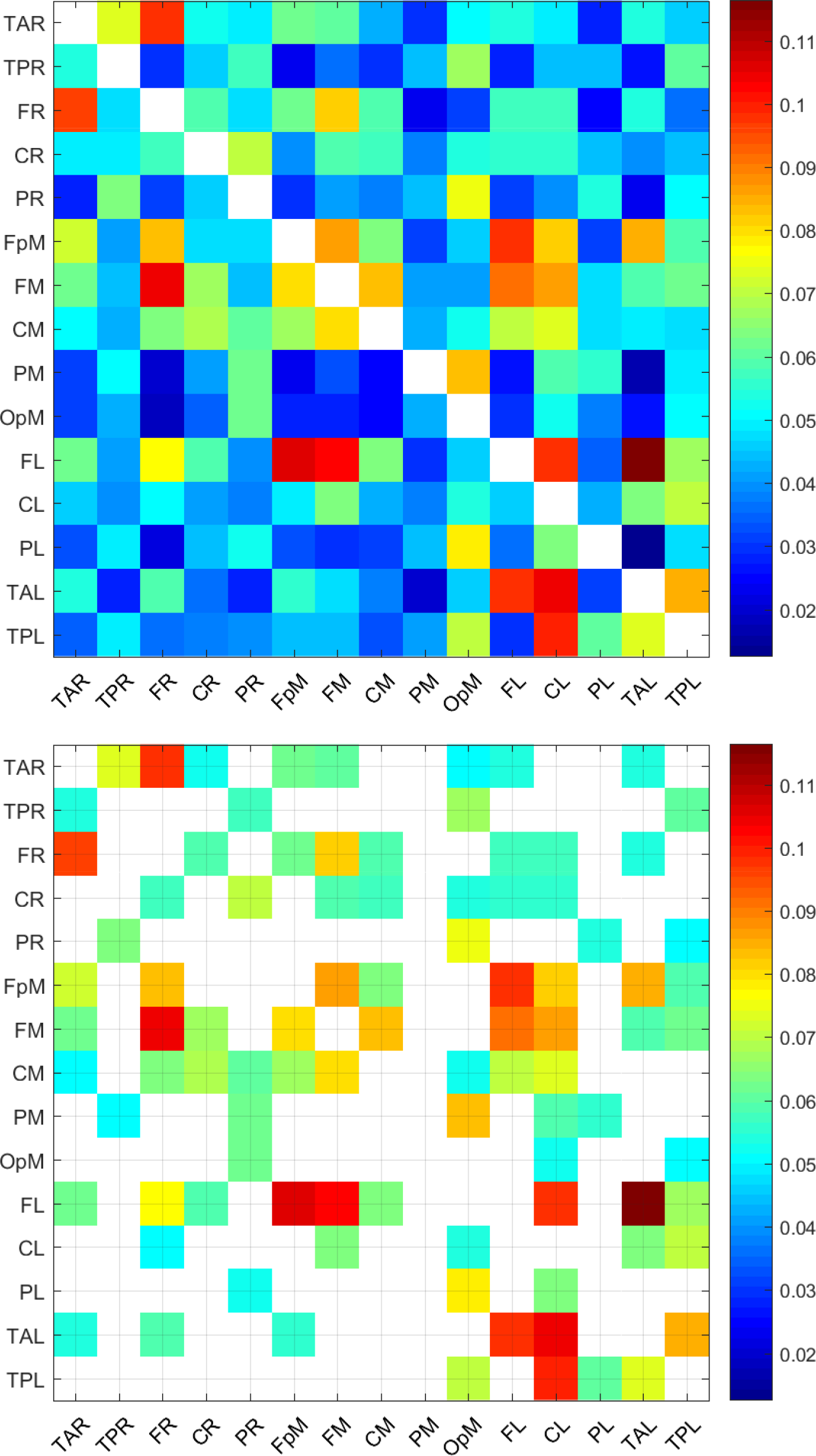
Mean normalized information flow rates, 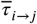, calculated over all the individuals in the study cohort (top) and corresponding values for connections above the threshold *τ*_*c*_ = 0.05 (bottom). There are 92 connections with 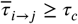.

A relevant question for interpreting the results is how many of the 210 connections (represented by the off-diagonal matrix cells) shown in *Figure 1* are important. As we discuss in Material and methods, it can be shown by *permutation testing* that the vast majority of the connections for all the individuals are statistically significant even at the *p* = 0.001 level. However, very low values of information flow rate, albeit statistically significant, imply that the relative impact of the *transmitter* series on the *receiver* is not neurologically important. On the other hand, there is no golden rule for selecting a threshold value above which connections are considered important (*Cohen, 2014*). Hereafter, we will consider that a connection *i* → *j* between two dipoles is *active in the ensemble sense* if the magnitude of the normalized information flow rate |*τ*_*i*→*j*_| exceeds the arbitrary threshold of *τ*_*c*_ = 0.05. This means that the entropic rate of change at the *receiver j* due to its interaction with the *transmitter* located at *i* is at least 5% of the total rate of entropy change at *j*. (We further comment on the selection *τ*_*c*_ = 0.05 in connection with *Figure 3* below.) The bottom panel of *Figure 1* shows the mean information flow rate for the connections that are active in the ensemble sense. The latter involve only connections such that 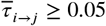. As evidenced in this plot, 92 out of the 210 inter-dipole pairs are connected on average, i.e., they exhibit 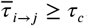.

The top thirty (30) active connections, ranked on the basis of 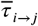, are listed in *Table 1* and displayed by arrows on an axial view schematic in *Figure 2*. All thirty connections correspond to positive values of 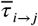 in the interval between 0.116 (highest) and 0.072 (lowest). All of them have values higher than the threshold *τ*_*c*_ = 0.05. Evaluating the thirty top connections (cf. *Figure 2*), the overall information flow pattern is predominantly left lateralized and consists of mostly short and medium range bidirectional connections linking the frontal, central and temporal regions of the brain. The possible neurological insights derived from *Table 1* and *Figure 2* are developed in the Discussion section.

**Table 1.** List of the most active connections based on the ensemble mean of the normalized information flow rate, *τ*_*i*→*j*_. All *τ*_*i*→*j*_ listed exceed the threshold *τ*_*c*_ = 0.05. The connections listed are included among those shown in the bottom matrix plot of *Figure 1*. The last column in the table reports the polarization which is equal to the sum of the signs of *τ*_*i*→*j*_ over the individuals as a percentage of the number of individuals in the study (*L* = 32) [see *Equation 6*].

| | From | To | Mean $\tau_{i \rightarrow j}$ | Polarization |
| --- | --- | --- | --- | --- |
| 1 | FL | TAL | 1.164062e-01 | 100 |
| 2 | FL | FpM | 1.062701e-01 | 100 |
| 3 | TAL | CL | 1.048785e-01 | 100 |
| 4 | FM | FR | 1.039752e-01 | 94 |
| 5 | FL | FM | 1.023856e-01 | 100 |
| 6 | TPL | CL | 9.902969e-02 | 100 |
| 7 | TAL | FL | 9.789913e-02 | 100 |
| 8 | TAR | FR | 9.752876e-02 | 100 |
| 9 | FpM | FL | 9.738423e-02 | 100 |
| 10 | FL | CL | 9.737341e-02 | 100 |
| 11 | FR | TAR | 9.643028e-02 | 100 |
| 12 | FM | FL | 9.108766e-02 | 100 |
| 13 | FpM | FM | 8.720060e-02 | 94 |
| 14 | FM | CL | 8.698357e-02 | 94 |
| 15 | TAL | TPL | 8.462667e-02 | 94 |
| 16 | FpM | TAL | 8.422089e-02 | 88 |
| 17 | PM | OpM | 8.336186e-02 | 100 |
| 18 | FpM | FR | 8.322882e-02 | 94 |
| 19 | FM | CM | 8.248654e-02 | 100 |
| 20 | FpM | CL | 8.198255e-02 | 100 |
| 21 | FR | FM | 8.095464e-02 | 94 |
| 22 | FM | FpM | 8.034528e-02 | 100 |
| 23 | CM | FM | 8.017243e-02 | 100 |
| 24 | PL | OpM | 7.770022e-02 | 100 |
| 25 | FL | FR | 7.733949e-02 | 100 |
| 26 | PR | OpM | 7.503928e-02 | 100 |
| 27 | CM | CL | 7.419229e-02 | 100 |
| 28 | TPL | TAL | 7.341683e-02 | 100 |
| 29 | TAR | TPR | 7.276059e-02 | 100 |
| 30 | FpM | TAR | 7.214605e-02 | 94 |

**Figure 2.**
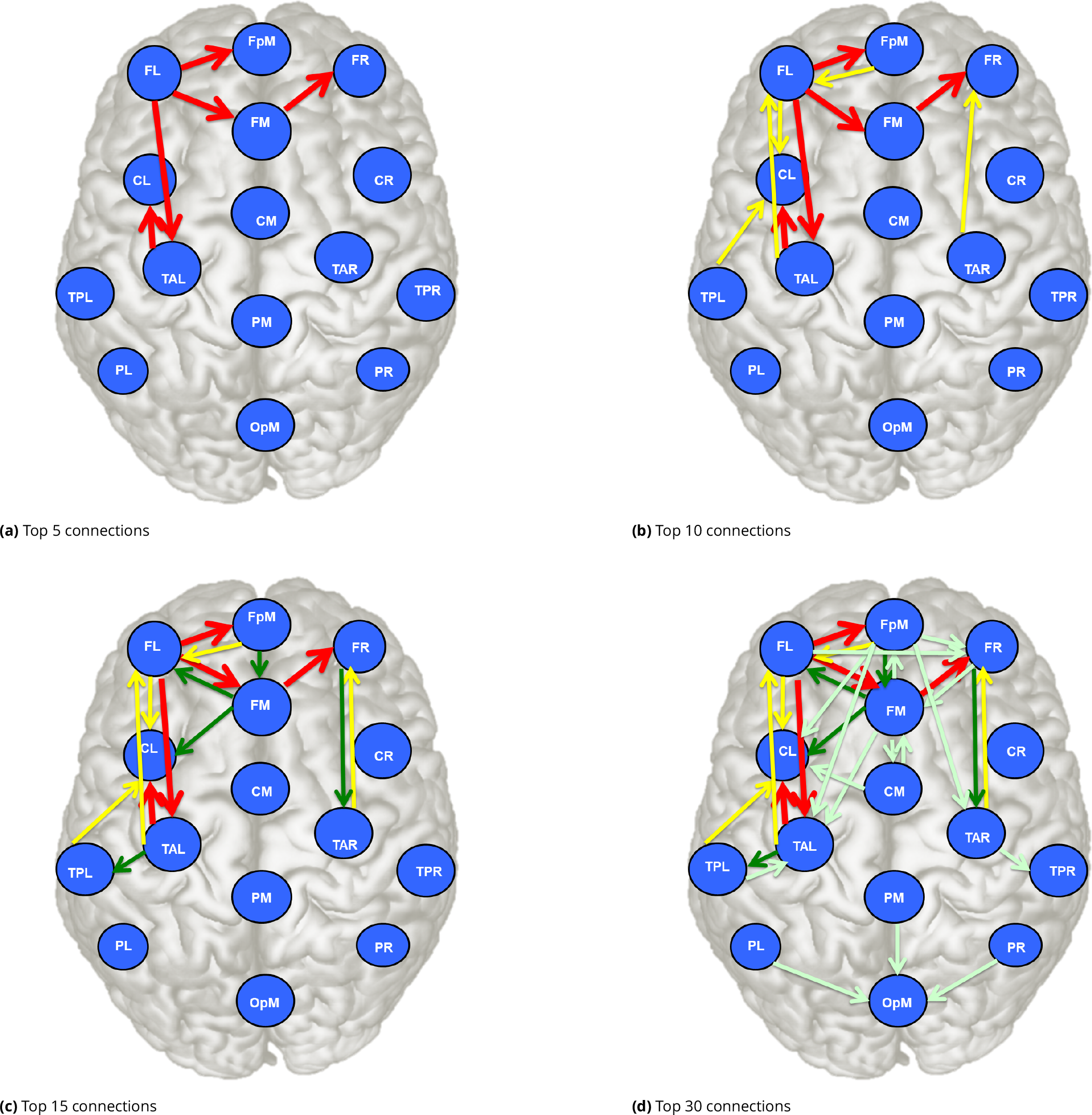
Axial-view schematic showing the main pathways of information flow based on the mean normalized information flow rates shown in *Table 1*. The schematic shows the locations of the sources in BESA’s 15 pre-defined regions (see Description of EEG data in Materials and methods. (a) Five most important connections shown in red arrows (color online). (b) Ten most important connections; connections ranked from six to ten are shown with yellow arrows. (c) Fifteen most important connections with those ranked from 11 to 15 shown with green arrows. (d) Thirty most important connections with those ranked from 15 to 30 shown in light green arrows (cf. *Table 1*).

The last column of *Table 1* displays the *polarization P* (*τ*_*i*→*j*_) of the information flow rate. This ensemble measure is given by the average sign of *τ*_*i*→*j*_ expressed as a percentage, i.e.,

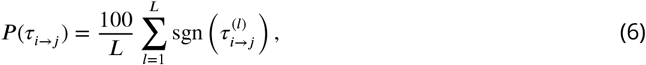

where sgn(⋅) is the sign function defined by sgn(*x*) = 1, if *x >* 0, sgn(*x*) = −1, if *x <* 0 and sgn(*x*) = 0, if *x* = 0. The polarization is a number close to ±100% if the sign of the *τ*_*i*→*j*_ is typically the same for all the individuals (if all the signs are the same the polarization is 100%). In *Table 1* the polarization varies between ≈ 88% and 100% and is less than 100% only for eight connections. This means that for the vast majority of connections, the variations between individuals affect the magnitude but not the sign of the normalized information flow rate.

A different way to view the relation between the ensemble mean 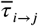 and the information flow rates of individuals is by counting for how many individuals each connection is active. Hereafter, we will consider that a connection *i* → *j* between two dipoles is *individually active* if the magnitude of the normalized information flow rate |*τ*_*i*→*j*_| exceeds the threshold *τ*_*c*_, which means that the percentage of the total entropy rate of the *receiver* due to its interaction with the *transmitter* is at least 5%. We assume that the threshold for individually active connections is the same as the threshold used for the ensemble mean of the information flow rate. However, this is not necessary in general.

We define the *frequency of activity*, *n*_*i*→*j*_(*τ*_*c*_), for the connection *i* → *j* as the number of individuals in the study cohort for which the specific connection is active. Hence,

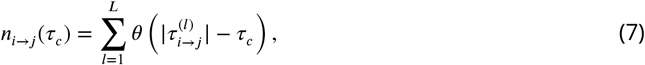

where *θ*(⋅) is the unit step function, i.e., *θ*(*x*) = 1, for *x* ≥ 0 and *θ*(*x*) = 0 for *x* < 0. The frequency of activity is evaluated over all the individuals and takes values integer *n*_*i*→*j*_(*τ*_*c*_) ∈ {0, 1, …, *L*}, where *L* = 32 is the number of individuals in the cohort. The frequency of activity depends on *τ*_*c*_, as higher values of *τ*_*c*_ imply a smaller number of active connections.

In *Figure 3*, we explore the correlation between the ensemble mean 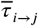 and the number of individually active connections *n*_*i*→*j*_(*τ*_*c*_). The scatter plot shows an almost linear dependence between the number of individually active connections and the respective value of 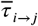 for values of 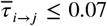 and appears to level off at higher values of 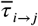. At the same time, the scatter also increases towards these higher values. We can also use this plot as a guide for selecting a suitable threshold for ensemble-based connectivity analysis, since it reveals how the threshold imposed on 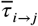 (shown as a vertical red line in *Figure 3*) affects the number of active connections, i.e., the number of markers to the right of the vertical line at *τ*_*c*_. However, note that the frequency of the connections in individuals (i.e., the values on the vertical axis) will change if a different threshold is used to estimate individual activity. Essentially, the plot would need to be redrawn for different values of individual *τ*_*c*_.

**Figure 3.**
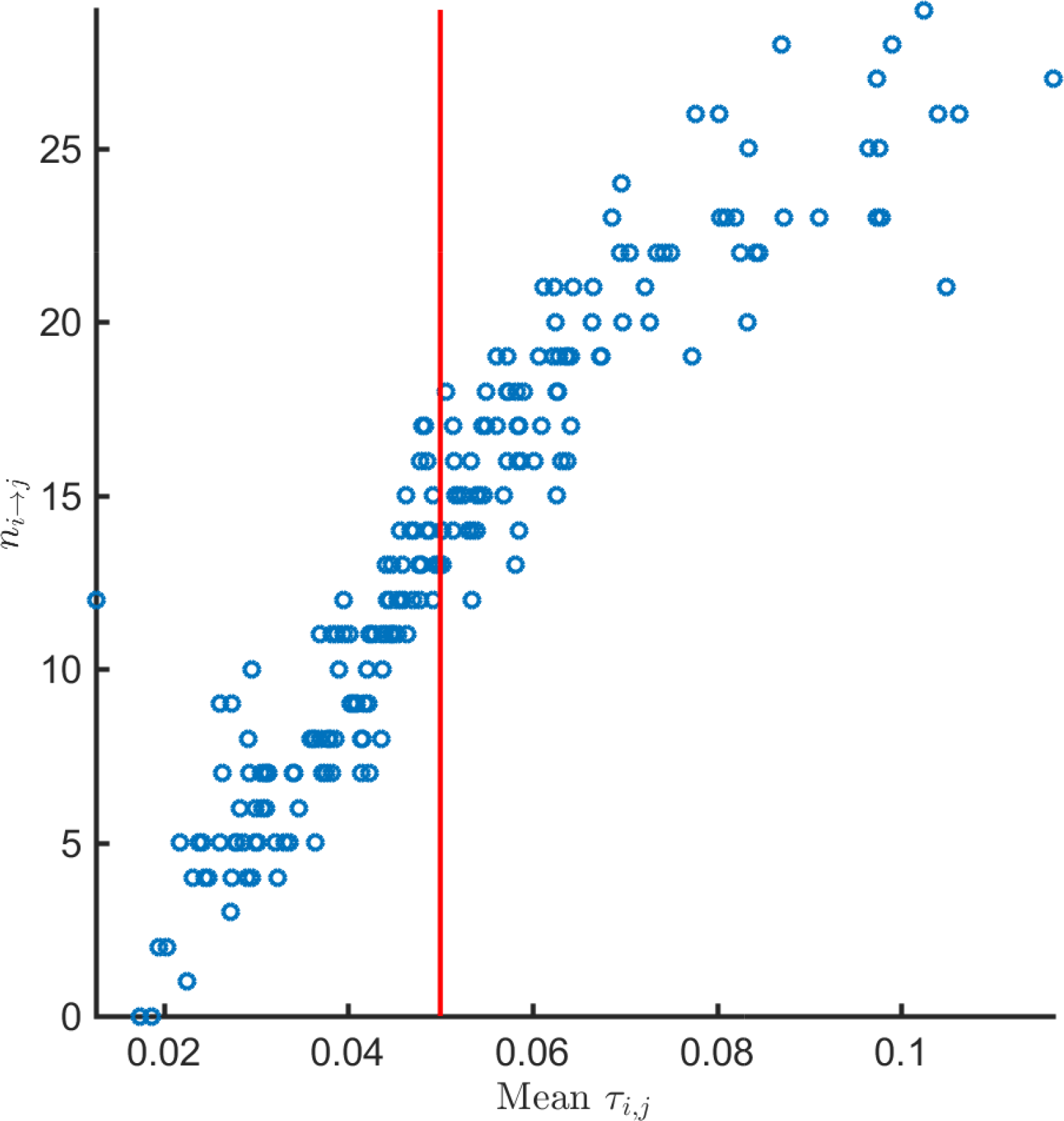
Frequency *n*_*i→j*_ (*τ_c_*) of individually active connections (based on the individual threshold *τ_c_* = 0.05) plotted against the ensemble mean of normalized flow rate coefficient *τ*_*i*→*j*_ for the connection *transmitter i* → *receiver j* for all *i* ≠ *j* = 1, …, 15. The plot comprises 210 points, each of which corresponds to a different *i* → *j* connection between two source dipoles. The frequency of activity *n*_*i→j*_ (*τ_c_*) is calculated over all *L* = 32 individuals in the cohort based on *Equation 7* and thus its upper bound is equal to *L*. The ensemble mean 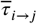 is calculated based on *Equation 5*. The vertical red line marks the threshold value *τ_c_* = 0.05. All the markers to the right of the vertical line correspond to connections *i* → *j* which are on average above the threshold.

### Information flow rate patterns per individual

To study the information flow across brain regions in individuals, we focus on the *individually active* source dipole pairs. As stated above, these are dipole pairs with *τ*_*i*→*j*_ whose magnitude (absolute value) exceeds the threshold *τ*_*c*_ = 0.05. We use the criterion |*τ*_*i→j*_| > *τ*_*c*_ instead of *τ*_*i*→*j*_ > *τ*_*c*_ since there are a few pairs (nine out of a total of 6720) with values of *τ*_*i*→*j*_ < −0.05.

For each of the 32 individuals in the study, we calculate 210 values of normalized inter-dipole information flow rates *τ*_*i*→*j*_. To calculate *τ*_*i*→*j*_, we use all the time points in the EEG time series. The *τ*_*i*→*j*_ values over all individuals range from −0.0794 to 0.3568. The matrix of the *τ*_*i*→*j*_ values *for each individual* is depicted in *Figure 4*-*Figure 7*. Each plot corresponds to a single individual and shows an *N*_*s*_ × *N*_*s*_ square grid that represents all the possible connections between sources. The value of each grid cell in *Figure 1* is equal to the average (evaluated over all the individuals) of the values of the respective grid cells in *Figure 4*-*Figure 7*.

**Figure 4.**
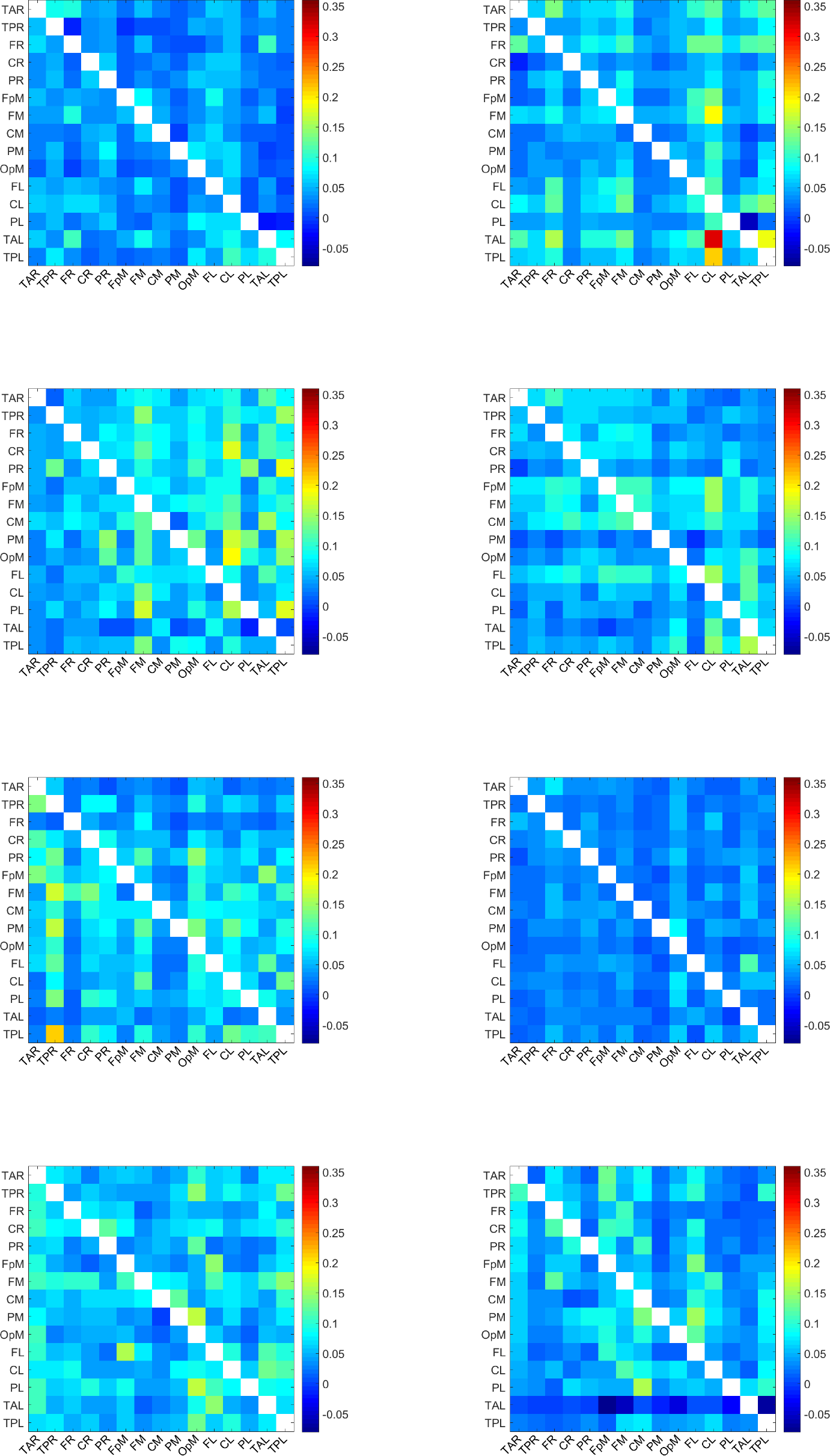
Map of *τ*_*i→j*_ values for the individuals 1-8. A uniform color map is used based on the range of *τ*_*i→j*_ values for all 32 individuals. The value of a grid cell (L1, L2), determined by the label L1 on the vertical axis and the label L2 on the horizontal axis, represents information flow from the dipole with label L1 to the dipole with label L2. The *τ*_*i→j*_ values over all dipoles and individuals range from −0.08 to 0.36.

**Figure 5.**
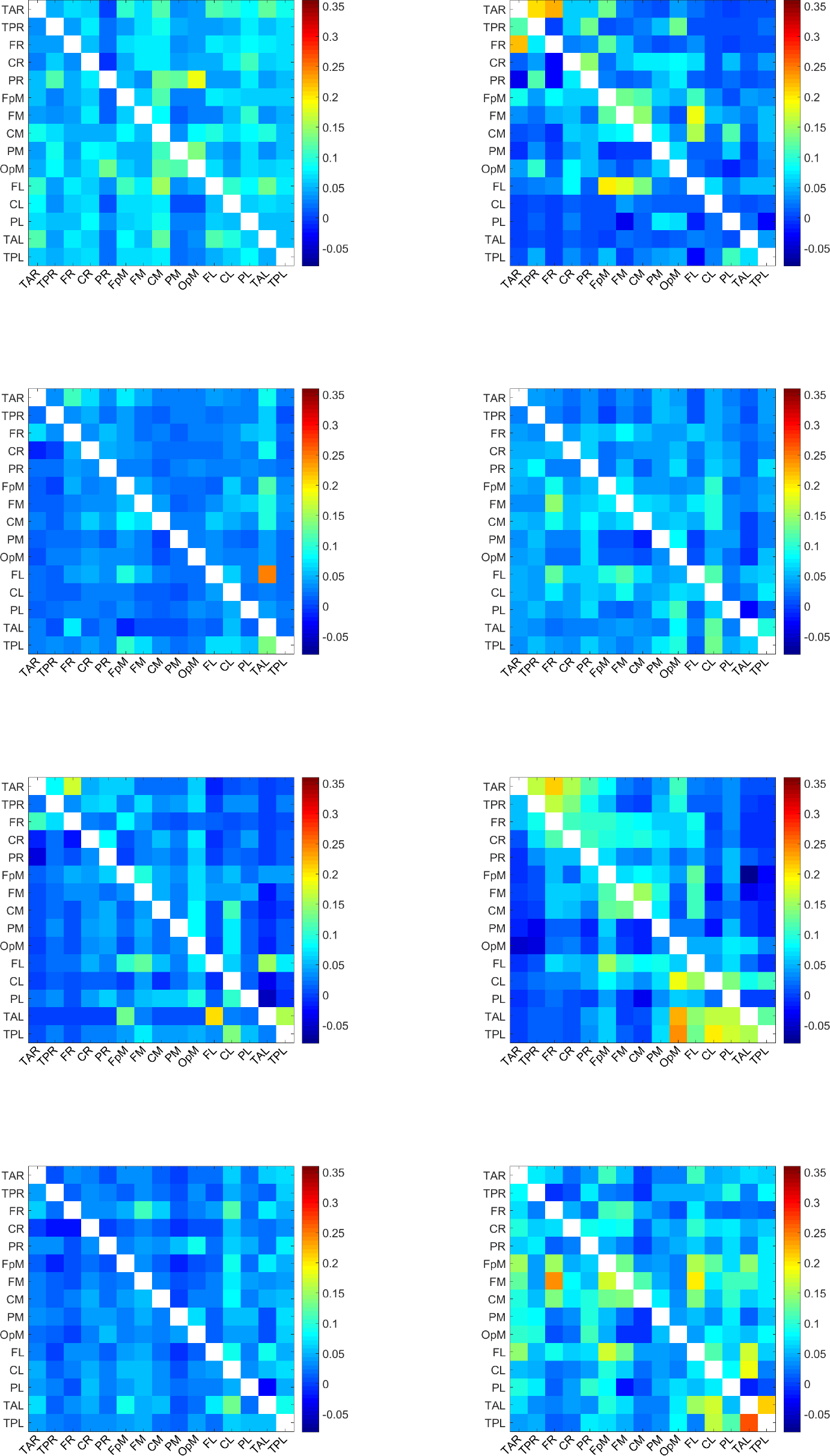
Map of *τ*_*i→j*_ values for the individuals 9-16. A uniform color map is used based on the range of *τ*_*i→j*_ values for all 32 individuals. The value of a grid cell (L1, L2), determined by the label L1 on the vertical axis and the label L2 on the horizontal axis, represents information flow from the dipole with label L1 to the dipole with label L2. The *τ*_*i→j*_ values over all dipoles and individuals range from −0.08 to 0.36.

**Figure 6.**
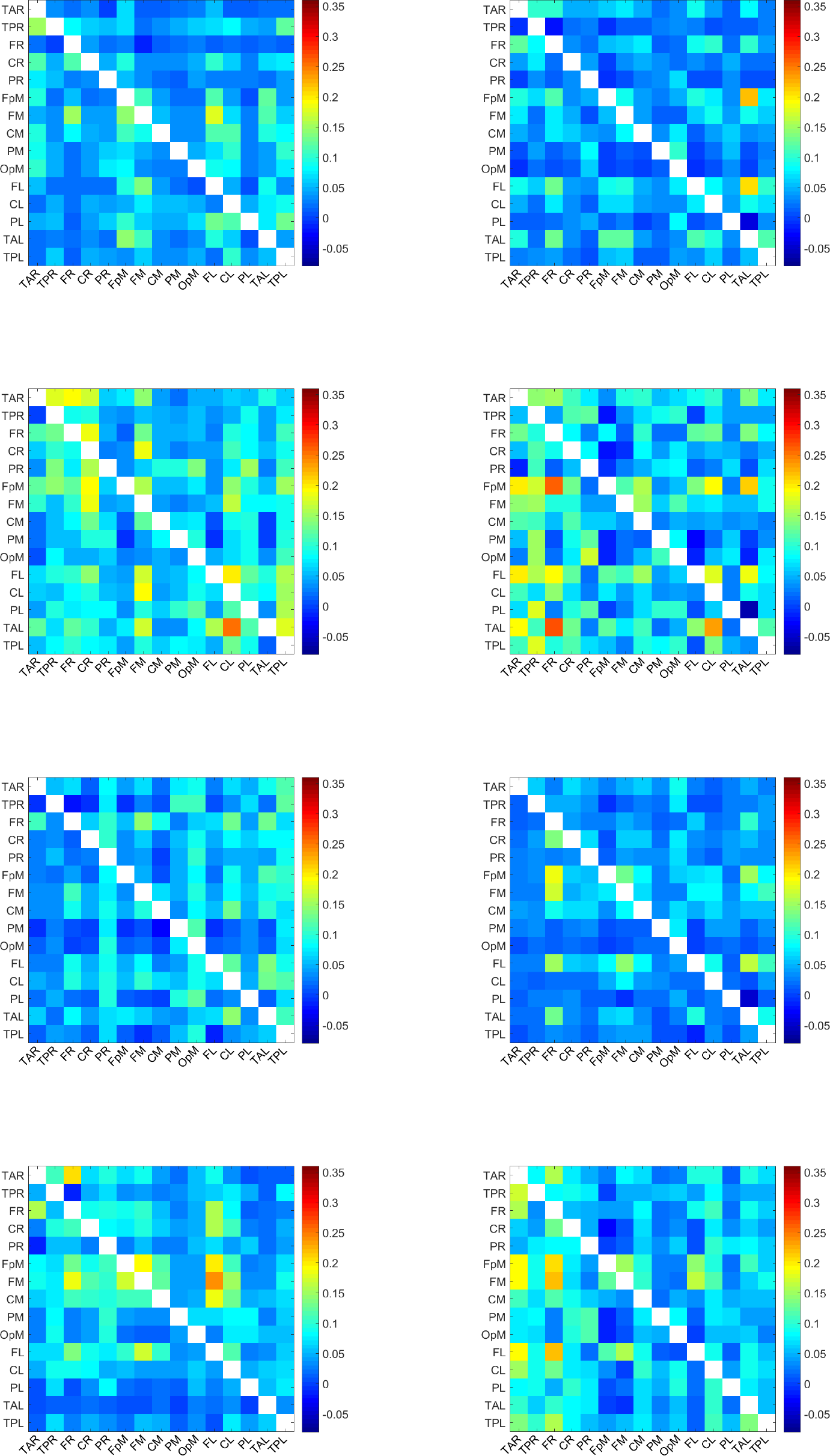
Map of *τ*_*i→j*_ values for the individuals 17-24. A uniform color map is used based on the range of *τ*_*i*→*j*_ values for all 32 individuals. The value of a grid cell (L1, L2), determined by the label L1 on the vertical axis and the label L2 on the horizontal axis, represents information flow from the dipole with label L1 to the dipole with label L2. The *τ_i_*_→*j*_ values over all dipoles and individuals range from −0.08 to 0.36.

**Figure 7.**
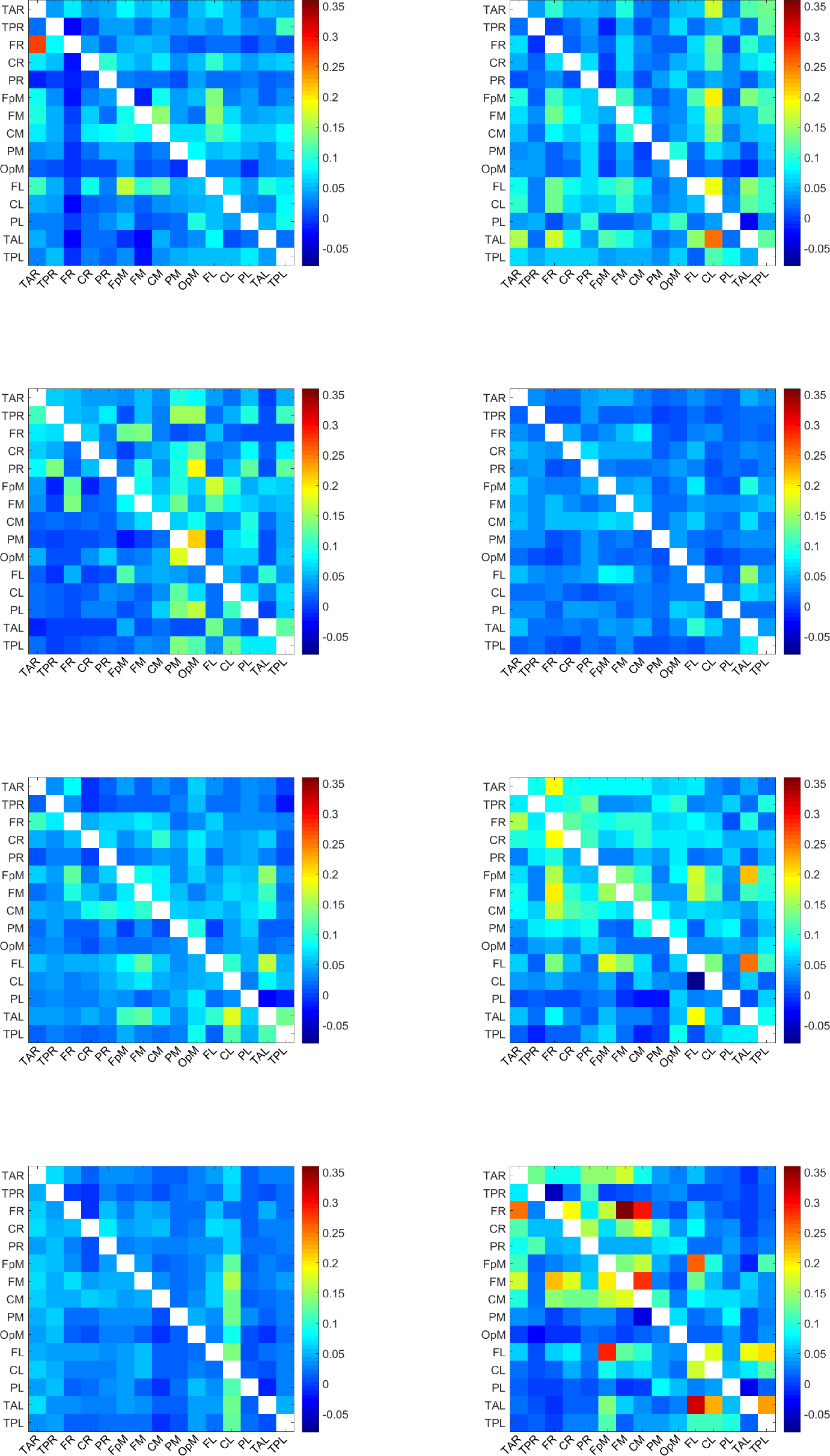
Map of *τ*_*i→j*_ values for the individuals 25-32. A uniform color map is used based on the range of *τ*_*i→j*_ values for all 32 individuals. The value of a grid cell (L1, L2), determined by the label L1 on the vertical axis and the label L2 on the horizontal axis, represents information flow from the dipole with label L1 to the dipole with label L2. The *τ*_*i→j*_ values over all dipoles and individuals range from −0.08 to 0.36.

These plots display the values of *τ*_*i*→*j*_ for all source dipole pairs, regardless of whether the connections are active with respect to the threshold *τ*_*c*_ or not. All the plots use a unified colormap based on the full *τ*_*i*→*j*_ range, i.e., [−0.08, 0.36] calculated over all dipole pairs and individuals. We note that the *τ*_*i*→*j*_ values are directional. For example, in the second plot (top right) of *Figure 4*, the cell labeled (TAL, CL) — near the bottom right of the grid — is colored red, which reflects a large value of *τ*_*i*→*j*_, while the cell marked by (CL, TAL) — above the main diagonal of the grid — has a much lower *τ*_*i*→*j*_. This indicates that the information flow from TAL has much higher impact on CL than the impact of CL on TAL.

The maximum *τ*_*i*→*j*_ observed among individuals is ≈ 0.36. This is about three times higher than the highest ensemble mean 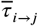 which is equal to 0.116 (cf. *Table 1*). This difference reflects the variability of the information flow rate values between individuals.

### Effective connectivity variations between individuals

To investigate the variability of the connectivity patterns between individuals, in *Figure 8* we plot the *frequency of activity*, *n*_*i→j*_(*τ*_*c*_), defined in *Equation 7*, for all *i* ≠ *j* = 1, …, 15 and for *τ*_*c*_ = 0.05. The main features evidenced in this plot are as follows:

1. Almost all the possible (208 out of 210) *transmitter i* → *receiver j* inter-dipole connections are active in at least one individual.
2. Of the 210 × 32 = 6720 pairs of inter-dipole connections that are available *in total* in the cohort of 32 individuals, only 2821 connections, or about 42% of the total number, are individually active (i.e., their magnitude is not less than *τ*_*c*_ = 0.05). This means that more than half of the current source dipole pairs in the study cohort are not strongly connected. In these pairs, the *transmitter* dipole does not strongly affect the *receiver* dipole.
3. In light of (1) and (2), we conclude that the active connections vary to some extent between individuals. For example, if the same set of about 42% connections were active for all (32) the individuals in the cohort, approximately 88 (i.e., 42% of 210) dipole pairs would be active. However, more than twice as many (i.e., 208) show active connections. In particular, 142 inter-dipole connections are active in ten or more individuals, forty inter-dipole connections (i.e., about 20% of the total connections) are active in twenty or more individuals, and twelve are active in more than 25 individuals.

**Figure 8.**
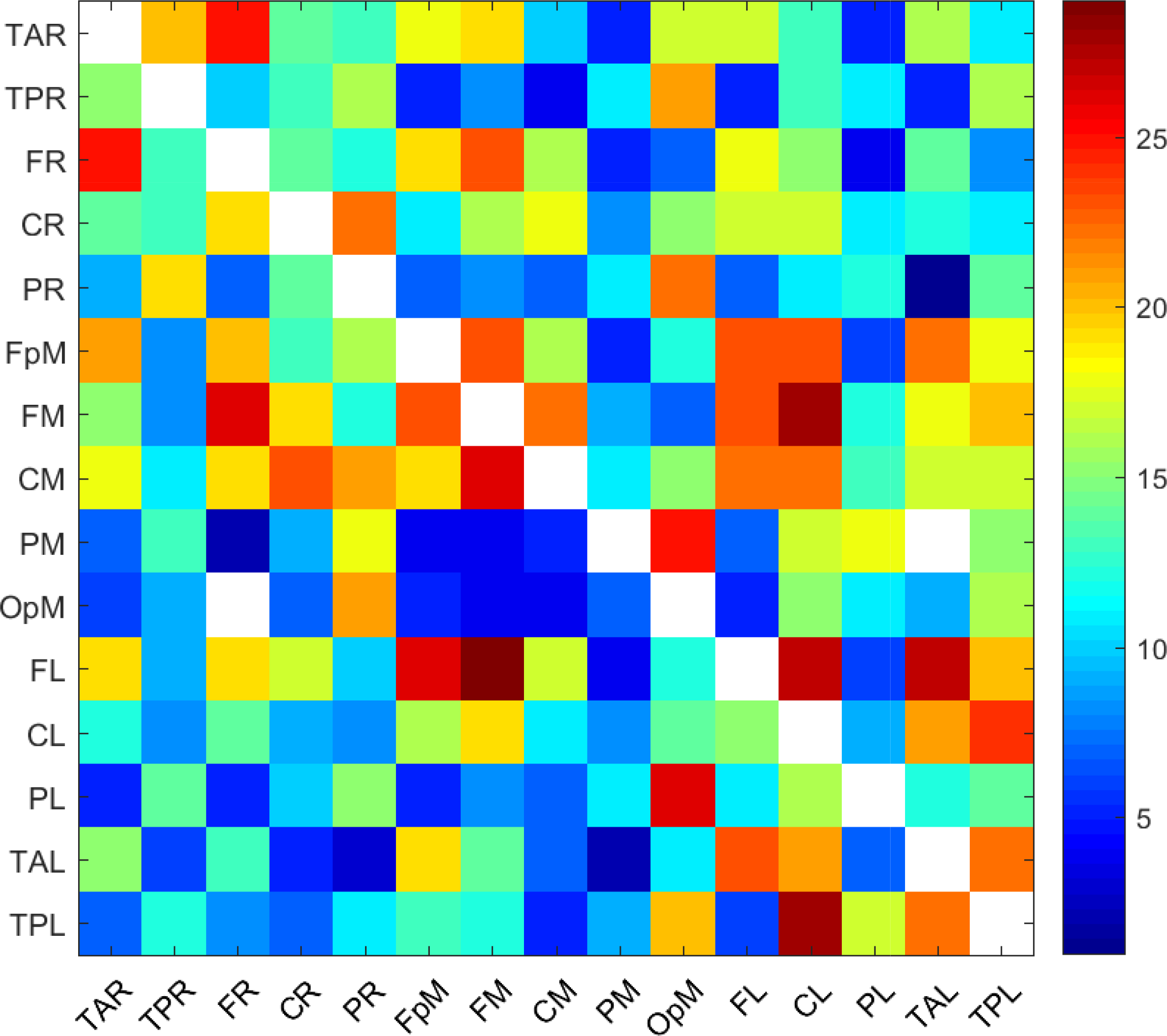
Frequency, *n*_*i→j*_(*τ*_*c*_), of active connections (i.e., source current dipole pairs with *τ*_*i→j*_ exceeding in magnitude the threshold value *τ_c_* = 0.05) calculated over all the individuals in the study. The value at each cell corresponds to the number of individuals for whom the specific connection is active. The white areas consist of cells with zero *n*_*i→j*_ (*τ*_*c*_), i.e., inactive connections. The diagonal cells are also white, since connections between the same *transmitter* and *receiver* are not meaningful.

The connectivity map in *Figure 8* exhibits a denser network of connections than the respective map in the bottom plot of *Figure 1*. The former shows the number of individuals for which a particular inter-dipole connection is individually active. Hence, it includes connections that are active in single individuals. On the other hand, the bottom plot in *Figure 1* displays the number of connections that are *active on average*, which is understandably smaller given the inter-subject variability. It is noteworthy that the five connections with the highest mean *τ*_*i*→*j*_, i.e., FL→TAL, FL→FpM, TAL→CL, FM→FR, FL→FM (cf. Figs. *Figure 1* and *Figure 2*, and *Table 1*), have relatively high frequencies of activity *n*_*i*→*j*_(0.05) = 27, 26, 21, 26, 29 respectively. In other words, these connections are active in most of the individuals (cf. *Figure 8*).

## Discussion

In this section we first discuss methodological aspects that are related to the information flow rate as well as its relation and differences with other connectivity measures. Then, we analyze the results that we obtained in this study in the context of the existing literature results on effective brain connectivity, focusing on the resting state of the adolescent brain.

### Brain connectivity measures and information flow

Functional measures of connectivity estimate non-directional relations and thus lead to undirected brain networks that fail to capture how one brain region influences another. However, such measures are still in common use (*Mill et al., 2017*). The simplest measure of functional connectivity is Pearson’s linear correlation coefficient (*Cohen, 2014*). Pearson’s coefficient fails to satisfactorily capture nonlinear dependence. Mutual information is a measure of functional connectivity that is based on information theory and can detect both linear and nonlinear relations (*Salvador et al., 2010*). Its calculation, however, requires the univariate probability distribution of each individual EEG time series, as well as the bivariate (joint) distribution for each pair of time series. Since long time series are required to estimate the bivariate distribution, the application of mutual information can be computationally intensive. Moreover, the method is sensitive to the number of bins used to estimate the probability histograms, and it fails to distinguish between nonlinear and linear, or positive and negative relations (*Cohen, 2014*).

On the other hand, measures of effective connectivity are directional variables which can distinguish the direction of information flow between brain regions. Measures of effective connectivity, such as *Granger causality* (*Kamiński et al., 2001*; *Hesse et al., 2003*; *Bressler and Seth, 2011*; *Seth et al., 2015*) and *transfer entropy* (*Schreiber, 2000*; *Liu and Aviyente, 2012*; *Salvador et al., 2010*; *Shovon et al., 2014*; *Hillebrand et al., 2016*), have been applied to EEG data to identify patterns of information flow in the functional brain networks during cognitive activity. Recently, *Muthuraman et al. (2015)* applied *renormalized partial directed coherence*, a measure based on the principle of Granger causality, to the combination of EEG and magnetoencephalography (MEG) signals to identify the direction of information flow between two signals and ultimately characterize the functional and effective connectivity in resting-state brain connectivity patterns. Thus, effective connectivity measures offer insights into the dynamics of the neuronal clusters that underpin cognitive function. Graphical models provide an intuitive tool for analyzing and visualizing associations and causal relationships and for modelling functional connectivity between brain regions (*Li and Wang, 2009*).

Granger causality analysis is based on the assumptions that (1) the time series are stationary, (2) interaction between the series can be described by means of a linear relation (typically a multivariate autoregressive model), (3) a specific model order can be defined, which determines how far in the past the coupling between two series extends, and (4) the innovation process of the linear model is described by Gaussian white noise (*Seth, 2007*; *Liu and Aviyente, 2012*; *Cohen, 2014*). This “plain vanilla” variety of Granger causality fails to detect nonlinear causal links (*Liu and Aviyente, 2012*; *Lin et al., 2017*). In such cases, nonlinear extensions of Granger causality are necessary (*Chen et al., 2004*; *Marinazzo et al., 2011*). However, such approaches are not yet conclusive since the selection of the degree of model nonlinearity and overfitting remain open issues (*Marinazzo et al., 2011*).

Transfer entropy is an extension of the concept of mutual information. It is based on the notion of relative entropy (also known as Kullback–Leibler divergence) and measures the difference between two probability distributions. For linear autoregressive systems driven by Gaussian white noise, Granger causality has been shown to be equivalent to transfer entropy (*Barnett et al., 2009*; *Liu and Aviyente, 2012*). Hence, the latter can be viewed as an extension of the former that can handle the dependence of non-Gaussian time series. Comparisons between Granger causality and transfer entropy are given in (*Bressler and Seth, 2011*; *Liu and Aviyente, 2012*). As stated above, Granger causality requires the specification of the order of the autoregressive processes involved. This model order, however, may depend on a number of variables including the conditions, the tasks executed (for task-oriented studies), and the EEG time series segments analyzed (*Cohen, 2014*). Transfer entropy makes fewer assumptions about the data than the standard Granger causality approach (*Vicente et al., 2011*). There are, nonetheless, challenges related to the calculation of transfer entropy, e.g., estimation by state-space partitioning, as discussed by *Bressler and Seth (2011)* and *Liang (2014)*.

Entropy and information content are key concepts in the definition of functional and effective brain connectivity measures (*Cohen, 2014*). In the thermodynamic sense, entropy is associated with disorder: a higher temperature implies higher entropy. In classical (as opposed to quantum mechanical) thermodynamics, the entropy ***S*** is calculated by means of the Gibbs formula *S* = −*k_B_ ∑_i_ p_i_* ln *p*_*i*_, where the summation is over the probabilities *p*_*i*_ of the system’s microstates (the index *i* should not to be confused with the location index of current source dipoles) and *k*_*B*_ is Boltzmann’s constant. In information theory, the entropy of a system with *N* states is defined in terms of Shannon’s formula 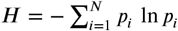, where *p*_*i*_, *i* = 1, …, *N* is the probability of the state indexed by *i*. If the natural logarithm is used in the definition (as was done above), Shannon entropy is measured in terms of *natural information units* (nats).

The Shannon entropy quantifies the unpredictability (uncertainty) of a stochastic system. High entropy implies that the result of a measurement is not only *a priori* unpredictable, but that the measurement itself provides new information which improves our knowledge of the system. On the other hand, low entropy means that the extant knowledge of the system allows us to predict quite well the outcome of the measurement and consequently, the measurement does not contain significant new information. Hence, higher entropy implies a higher level of unpredictability, while lower entropy implies that efficient, “compressed” representations are possible (i.e., a parameter set of lower dimensionality can be used to represent the system). In complex systems, there are interactions between different components. We can intuitively view information flow from component *X* to component *Y* as the amount of the uncertainty of *Y* that is resolved by the past states of *X*. If the past states of the component *X* do not affect the current state of *Y*, there is no information flow from *X* to *Y* (*Bossomaier et al., 2016*). On the other hand, the past states of *X* may reduce or improve the predictability of *Y*, thus implying information flow from *X* to *Y*. Currently used measures of functional and effective brain connectivity are based on the concept of the absolute Shannon entropy. The concept of Shannon entropy has been generalized to dynamical systems that are not necessarily stochastic, by means of the Kolmogorov-Sinai entropy (*Gutzwiller, 1990*) which quantifies the unpredictability of future states of the system.

The *Liang-Kleeman* information flow rate is a recently developed measure which is also based on the concept of Shannon entropy (*Liang, 2008*, 2013b,a, 2014). However, the information flow formalism can be derived using either absolute or relative entropy. In two dimensions (i.e., for a system of two time series) this was shown by Liang (2013b, 2014). *Relative entropy* (Kullback–Leibler divergence) measures how much information is added to a given system with respect to the information contained in the initial probability distribution. Recently, *Liang (2018)* has shown that the relative entropy formulation of the information flow rate is also valid for stochastic dynamical systems with *N >* 2 dimensions (i.e., systems involving *N* potentially coupled time series, where *N* is an arbitrary integer value).

The information flow rate formulation is based on the theory of dynamical systems, in contrast with transfer entropy which is a statistically motivated measure of information transfer. The information flow rate aims to address the computational shortcomings of transfer entropy (requirement for long time series, computational complexity, estimation of bivariate probability distribution) as well as spurious causal associations (*Liang, 2016*). The information flow rate provides an easy-to-compute *directional (asymmetric)* measure of dependence between pairs of time series that can be evaluated from a single realization of each series and does not require the estimation of transition probabilities. Unlike Granger causality, the information flow rate concept does not require a specific model structure, Gaussian statistics, or stationarity (*Liang, 2015*) and can also be applied to deterministic nonlinear systems (*Liang, 2016*). These could be important advantages of information flow, since the EEG signals exhibit non-stationary features evidenced in transitions between quasi-stationary periods and nonlinear dynamic behavior (*Blanco et al., 1995*; *Kaplan et al., 2005*; *Klonowski, 2009*), while the correct model structure is never known *a priori*. However, further study is needed to compare in detail the performance of the information flow rate against the standard methods of assessing brain connectivity.

Finally, we draw attention to an ongoing discussion in the literature regarding the very definition of *effective connectivity*, e.g. (*Lindquist, 2008*). Friston argues that effective connectivity should be based on dynamic models, such as the Dynamic Causal Models (DCMs) (*Friston, 2011*). Model-based connectivity methods assume a well-defined biophysical model of neuronal dynamics (*Sakkalis, 2011*). Friston also opines that data-driven models, such as Granger causality, provide functional connectivity measures, a view that is echoed by *Bastos and Schoffelen (2015)*. On the other hand, several other publications referenced in this paper, including the reviews (*Sakkalis, 2011*; *Bastos and Schoffelen, 2015*), refer to causality-based methods, including data-driven methods such as Granger causality and transfer entropy, as effective connectivity measures. We follow the latter viewpoint, according to which the information flow rate is an effective connectivity measure.

### Comparison of brain connectivity results with literature

In recent years, a number of advances facilitating the study of functional connectivity of the brain have thoroughly transformed our understanding of the activity present in the brain in absence of “any imposed stimuli, task performance or other behaviourally salient events” [for a review, see (Snyder and Raichle, 2012)]. This “resting state” of the brain is characterized by spontaneous, coherent fluctuations of blood-oxygen-level-dependent (BOLD) as well as electromagnetic signals from functionally distinct brain regions. fMRI studies were the first to show that subsets of these regions tend to act in concert, giving rise to functionally relevant “resting-state” brain networks (*Raichle et al., 2001*; *Greicius et al., 2009*) that provide a basis for information processing and coordinated activity. More recently, *Yuan et al. (2016)* and *Liu et al. (2017)* have found that functional resting-state networks can also be extracted from source-space EEG data, and *Hillebrand et al. (2016)* have done the same using MEG data. The most commonly reported resting-state functional networks observed in children (*Muetzel et al., 2016*), adolescents (*Borich et al., 2015*) and adults (*Yuan et al., 2016*; *Liu et al., 2017*) (and references therein) include the visual, the fronto-parietal, the sensory motor and the default mode network (DMN). These studies also highlight that the above resting-state functional networks are not independent, and that there is a high degree of interconnections between them. To date, however, very few studies have investigated the information flow between the different networks.

To our knowledge, there are only three studies that have investigated the source space information flow pathways in adults (age: over 20 years) during eyes closed resting state, and considered the relationship between these pathways and the underlying functional networks: (1) *Michels et al. (2013)* study EEG data using the partial directed coherence (PDC) measure, which is based on Granger causality, to quantify effective connectivity; (2) *Muthuraman et al. (2015)* analyze both EEG and MEG data, also by means of the partial directed coherence (PDC) measure; and (3) *Hillebrand et al. (2016)* study MEG recordings using directed phase transfer entropy (dPTE) to assess effective connectivity. All three studies find that the dominant pattern in adults is a posterior to anterior flow, originating in the regions associated with the primary visual cortex and the posterior DMN, and flowing to the frontal regions. *Michels et al. (2013)* and *Muthuraman et al. (2015)* observe only one-way connectivity between the brain regions. However, *Hillebrand et al. (2016)* find that the dominant patterns are complemented by weaker anterior to posterior connections which make the flow bidirectional at finer connection strength resolution.

Only *Michels et al. (2013)* have investigated source-space resting-state directed connectivity in children (mean age: 10 years). They find that the dominant flow pattern is opposite to that observed in the adults, with activation originating in the anterior (i.e. pre-frontal) regions and terminating in the posterior (parietal/occipital) regions. One possible explanation is that the anterior to posterior flow in children indicates modulation of lower-order sensory-motor information from frontal regions (*Emberson et al., 2015*; *Taylor and Khan, 2000*). Admittedly, there are obvious gaps in our understanding of the resting-state dynamics over the course of development. More studies of the resting-state dynamics in children, as well as detailed comparisons with other populations at different stages of development are needed to fully contextualize these findings.

The present study provides the first critical step towards understanding information flow in the brain during a key transition stage between childhood and adulthood. We have analyzed resting-state EEG data from an intermediate population, a cohort of adolescents (mean age: 16 years). Using the Liang-Kleeman information flow rate as a measure of effective brain connectivity, we find that of the 30 *active* connections in adolescent brains (based on the ensemble means of the normalized information flow rate), the five strongest (cf. red arrows in *Figure 2a*) mostly originate in the left frontal region of the brain and flow to left temporal and mid-frontal regions. Including the next ten connections (cf. yellow and dark green arrows in Figs. *Figure 2b* and *Figure 2c*) extends the active areas of the brain beyond the frontal region to encompass adjacent posterior regions (i.e., central and temporal), with the information flow pattern becoming largely bidirectional but still strongly left lateralized. The final fifteen connections (cf. light green arrows in *Figure 2d*) are characterized by information flows between mainly the left and mid anterior regions. They also show some slightly lower level activity on the right side of the brain, an indication of inter-hemispheric flow between the left and right frontal regions, and the emergence of connections in the posterior regions (i.e. parietal to occipital). Overall, the information flow pattern suggested by the thirty connections is highly left lateralized and comprises mostly short and medium range bidirectional connections that link the frontal, central and temporal regions of the brain.

The above results are reminiscent of the basic directed connectivity pattern observed by *Michels et al. (2013)* in young children but with one important difference. In early adolescence, the pattern of information flow manifests an additional layer of complexity indicated by bidirectional communication between brain regions that *Hillebrand et al. (2016)* observe in the adults and which they interpret as feedback loops. In effect, the pattern that we observe in our cohort suggests a progression towards maturation of the adolescent brain.

Similarly, the lateralization of the information flow we observe is also a reflection of an earlier developmental stage. *Agcaoglu et al. (2015)* studied individuals ranging from 12 to 71 years and observed that the resting-state networks of young individuals are highly lateralized, with the default mode network, attention and frontal networks being strongly left lateralized. With age, however, this lateralization decreases and the network becomes more symmetric. In fact, the degree of interaction between networks, the order in which the networks are activated, the organization and the strength of the interactions within individual networks (including the extent to which they are lateralized), all change over development (*Muetzel et al., 2016*).

The fact that both functional and effective connectivity changes as the brain matures is not entirely surprising. It is well known that the brain undergoes considerable structural changes during the transition from puberty to adulthood (*Shaw et al., 2008*) as manifested by significant increase (decrease) in the volume of white (grey) matter (*Gogtay et al., 2004*; *Paus, 2005*; *Toga et al., 2006*; *Lebel and Beaulieu, 2011*). For example, *Lebel and Beaulieu (2011)* have shown that while the maturation of the projection fibers linking the primary sensorimotor cortical regions with lower-order subcortical sensory areas and the commissural fibers connecting the two hemispheres of the brain is mostly complete by late adolescence, the maturation of the association tracts, particularly the superior longitudinal and fronto-occipital fasciculi that connect the occipital and the frontal regions of the brain, continues well into the twenties. Functionally, these long association fibers are correlated with increasing long-range EEG coherence and synchronization (*Miskovic et al., 2015*).

Finally, we have also identified significant variability of effective connectivity between individuals based on the patterns of information flow rate between brain regions. We have presented and discussed graphical tools for visualizing and characterizing variability between individuals including dipole-dipole connectivity plots that account for all the individuals in the cohort, e.g., *Figure 8*. The variability of the brain’s resting-state functional and effective connectivity across individuals and over time are topics of considerable interest within both research and clinical settings. *Hutchison et al. (2013)* and *Hirayama et al. (2016)* (see also references therein) argue that the variability of the connectivity matrix between individuals is not due to noise but is associated with individual variances in mental/vigilance states and cognitive function. They also note that there are reports of the temporal dynamics of the connectivity matrix being affected by brain health, which raises the exciting possibility that, in the future, the associated features could serve as disease/injury biomarkers. The significant advantages of the new data-driven measure of effective brain connectivity discussed in this paper (i.e., ease of calculation, sensitivity to both linear and nonlinear relations, independence from a specific model structure and the stationarity assumptions), make it especially well suited for exploring these exciting new directions.

## Materials and Methods

In this section we briefly describe the EEG dataset. We then present the Liang-Kleeman directional information flow rate that will be used for the analysis of resting-state EEG brain connectivity. We also discuss how to numerically calculate and evaluate the statistical significance of the information flow rate obtained from the EEG data.

### Ethics Statement

This study was approved by the University of British Columbia Clinical Research Ethics Board (Approval number: H17-02973). The adolescents’ parents gave written informed consent for their children’s participation under the approval of the ethics committee of the University of British Columbia and in accordance with the Helsinki declaration. All participants provided assent.

### Participants

Thirty-two (32) right-handed male adolescents (mean age: 15.8 yrs; SD: ±1.3) participated in this study. Exclusion criteria for all individuals included focal neurologic deficits, pathology and/or those on prescription medications for neurological or psychiatric conditions. Parents signed an informed consent form that was approved by the University of British Columbia and all participants provided assent.

### Description of EEG data

Between 5−8 minutes of resting-state EEG data were collected while participants had their eyes closed, using a 64-channel Hydrogel Geodesic SensorNet (EGI, Eugene, OR) connected to a Net Amps 300 amplifier (*Virji-Babul et al., 2014*). The sensor-space signals were referenced to the vertex (Cz) and recorded at a sampling rate of *f*_*s*_ = 250 Hz. The scalp electrode impedance values were typically less than 50 *k*Ω. To eliminate artifacts associated with attaching (removing) the cap, 750 data points were removed from the beginning (end) of each time series. (This corresponds to removing data with a total duration of 6s.) The EEG time series were then filtered using a band-pass filter (4–50 Hz) and a notch filter (60 Hz), as described in (*Porter et al., 2017*) [see also (*Rotem-Kohavi et al., 2014*, *2017*)], to remove signal drift and line noise. In addition, Independent Component Analysis (ICA) was used to identify, decompose and remove eye blinks. Finally, the data were visually inspected and epochs with motion as well as additional ocular artifacts were excluded, as were channels with excessive noise. Each of the resulting EEG series used in this study involves between 67,845 and 114,304 time points.

Next, we used the Brain Electrical Analysis (BESA) Version 6.3 software^1^ (MEGIS Software GmbH, Gräfelfing, Germany) to map the cleaned sensor-space data to source waveforms. The voltages from the available sensor channels were first interpolated to voltages at 81 predefined scalp locations that comprise BESA’s Standard-81 10-10 Virtual Montage (*BESA Wiki, 2018*) and re-referenced to the average reference by subtracting the mean voltage of the full set of 81 virtual scalp electrodes. BESA uses spherical splines interpolation to perform this mapping (*Perrin et al., 1989*; *Scherg et al., 2002*). The interpolation offers a consistent way of dealing with occasional bad channels while maintaining a common montage across all the individuals. Thereafter, we use the BESA montage method (*Scherg et al., 2002*) to compute source waveforms. Since resting-state activity is not localized, we used the BR_Brain Regions montage which is derived from 15 pre-defined regional sources that are symmetrically distributed over the entire brain. The respective brain regions involved in this montage are listed in Table 2 and shown in *Figure 2*. BESA uses a linear inverse operator of the lead field matrix, which accounts for the topography of the sources included in the BR_Brain Regions montage, to calculate the source waveforms (*Scherg et al., 2002*). The composite source activity in each brain region is represented by a single regional source. Each source is modeled as a current dipole whose moment is specified in terms of a *local* orthogonal coordinate system with basis vectors commonly labelled as radial (r), horizontal (h), and vertical (v). Thus, the source waveforms represent time series of the fifteen current dipoles. Finally, the resulting data were exported to MATLAB for the analysis described below.

**Table 2.** List of the 15 brain regions used in EEG source space reconstruction in BESA.

| Source Dipole Label | Brain Region |
| --- | --- |
| FL | Frontal, left hemisphere |
| FpM | Fronto-polar midline |
| FR | Frontal, right hemisphere |
| FM | Frontal midline |
| CL | Central, left hemisphere |
| CM | Central midline |
| CR | Central, right hemisphere |
| TPL | Posterior temporal, left hemisphere |
| TAL | Anterior temporal, left hemisphere |
| TAR | Anterior temporal, right hemisphere |
| TPR | Posterior temporal, right hemisphere |
| PM | Parietal midline |
| PL | Parietal, left hemisphere |
| PR | Parietal, right hemisphere |
| OpM | Occipital-polar midline |

### Definition of inter-dipole information flow rate

In the following, 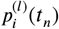 will denote the time series quantifying the time-varying strength (magnitude) of the current dipole moment at the source location *i* (where *i* = 1, …, *N_s_* = 15) for the individual indexed by *l* (where *l* = 1, …, *L* = 32), at time *t*_*n*_ = *n* Δ*t*, where *n* = 1, …, *N* is the time index and Δ*t* = 4ms is the *time step*. In terms of the dipole moment components 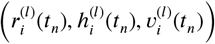 in the local (*r, v, h*) system, the magnitude of the dipole moment is given by

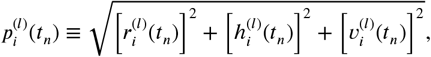

For completeness, we note that in general both the strength and the orientation of the current dipoles vary with time; however, in the present study, we track only their strength. In addition, we drop the individual index *l* if there is no risk of confusion. For short, we will also write *p_i,n_* = *p*_*i*_(*t*_*n*_).

Unlike the Pearson correlation coefficient which satisfies 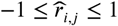 due to Schwartz’s inequality, the magnitude of the coefficient 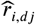 is not constrained to be less than one. This is due to the normalization of 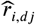 by the standard deviation of *p*_*j*_ instead of the standard deviation of the temporal derivative 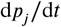 (cf. *Equation 3*).

Based on the discussion of Shannon entropy (cf. Discussion section), a positive (negative) rate of information flow from *i* → *j* (*T*_*i*→*j*_) indicates that the interaction between the two series leads to an increase (decrease) in the entropy of the series *p*_*j*_. Equivalently, it signifies that the *receiver* series becomes more (less) unpredictable due to its interaction with the *transmitter* series. The predictability of each time series is negatively correlated with the entropy.

While the information flow rate coefficients *T*_*i*→*j*_ were initially formulated for bi-variate systems that involve two interacting time series, Liang has recently proved theoretically that the equations above are also valid for *N*-variate, deterministic or stochastic systems (*Liang, 2016*, *2018*). In addition, even though the estimator of *T*_*i*→*j*_ has been derived using the assumption of a linear system, it has been successfully applied to identify causal connections in nonlinear systems as well (*Liang, 2014*, *2016*).

### Normalized information flow rate

The information flow rate is based on the notion of information entropy. A positive *T*_2→1_ implies that the *transmitter* series *p*_2_ increases the entropy of the *receiver p*_1_, while a negative *T*_2→1_ implies the opposite. By comparing *T*_*i*→*j*_ with *T*_*j*→*i*_(in the latter the roles of *transmitter* and *receiver* are reversed), we can determine which series transfers more information to the other series. However, this comparison does not reveal which of the two series is affected more due to its interaction with the other, because the coefficient does not account for the entropy change of each series due to the intrinsic evolution and possible stochastic effects. In order to quantify the impact of the entropy transferred to the *receiver* from a *transmitter* series, we need to know the extent to which the information transfer affects the predictability of the *receiver*, relative to all the other influences acting on the *receiver*.

The total rate of entropy change of *p*_*j*_ (*receiver*) depends not only on the information flow from *p*_*i*_(*transmitter*), which is determined by the rate *T*_*i*→*j*_, but also on 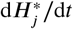 and 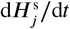. The term 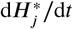(*intrinsic entropy rate*) represents the entropy rate of change due to the change of the phase space in the direction *p*_*j*_. The term 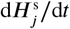(*noise-induced entropy rate*) represents the impact of stochastic effects in the dynamical system that underlies the evolution of *p*_*j*_ (*Liang, 2008*). Hence, as proposed by *Liang (2015)*, a suitable normalization factor for the information flow rate from *p*_*i*_ to *p*_*j*_ is derived by adding the absolute values of the three rates that contribute to the total rate of entropy change of the *receiver p_j_*, i.e.,

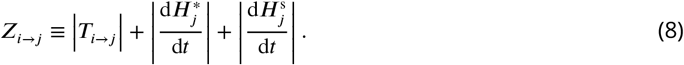

Based on *Equation 8*, *Z*_*i*→*j*_ is a non-negative number bounded from below by |*T*_*i*→*j*_|. In addition, *Z*_*i*→*j*_ cannot be zero unless the rate of change of the intrinsic and stochastic entropy components are zero. This can only happen if *p*_*j*_ is constant in time, which is not relevant for the EEG time series. Thus, the *normalized information flow rate* from the *transmitter p_i_* to the *receiver p_j_* is defined as (*Liang, 2015*)

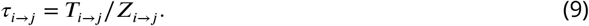

According to *Equation 9*, *τ*_*i*→*j*_ measures the percentage of the total entropy rate of change for *p*_*j*_ which is due to its interaction with *p*_*i*_. The calculation of the terms which contribute to *Z*_*i*→*j*_ from the data is explained in Appendix 1.

Based on the above analysis, the normalized information flow rate, *τ*_*i*→*j*_, has several advantages over the un-normalized coefficient, *T*_*i*→*j*_, the most important being that (1) *τ*_*i*→*j*_ does not explicitly depend on the finite difference step (item 3 in Box 4), and (2) it measures the importance of information flow from the *transmitter* to the *receiver* (items 4 and 5 in Box 4). Hence, *τ*_*i*→*j*_ is a suitable measure for investigating patterns of information flow between different regions of the brain and therefore for assessing effective connectivity.

#### Box 3. The main properties of the information flow rate *T*_*i*→*j*_ are as follows

1. In general, the correlation coefficients between 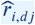 are not symmetric under interchange of *i* and *j*, i.e.,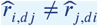. The asymmetry of 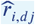 with respect to the interchange of *i* and *j* introduces directionality in the information flow rate coefficients, which implies that, in general, 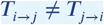.
2. For *i* = *j*, in light of 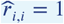 both the numerator and the denominator on the right-hand side of *Equation 1* become zero. Thus, *T*_*i→i*_ is undetermined; however, this is not an issue, because the quantities of interest are the rates of information flow between different time series.
3. The presence of the coefficients 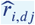 and 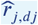 in the numerator on the right-hand side of *Equation 1* implies that *T*_*i*→*j*_ is proportional to the inverse of the finite difference time step, i.e., ∝ 1/*kΔt*, where *kΔt* is the time step used to calculate the time derivative (cf. *Equation 4*.)

#### Box 4. The main properties of the *normalized information flow rate τ*_*i→j*_ are as follows

1. The coefficient *τ_*i→j*_ is, in general, asymmetric, i.e., *τ*_*i*→*j*_* ≠ *τ*_*j*→*i*_ for *i* ≠ *j*.
2. The *τ*_*i*→*j*_ can take negative or positive values with magnitude less than one, i.e., −1 ≤ *τ_i,j_* ≤ 1. Positive values of *τ*_*i*→*j*_ imply that the *transmitter p_i_* tends to increase the entropy of the *receiver p_j_* (i.e., it increases its uncertainty), while negative values imply that *p*_*i*_ reduces the entropy of *p*_*j*_.
3. The *τ*_*i*→*j*_ does not explicitly depend on the finite difference step *kΔt*. This is due to the fact that both the numerator and the denominator in *Equation 9* are proportional to 1/*kΔt*.
4. The *τ*_*i*→*j*_ measures the relative importance of the entropy change in the *receiver* series *p*_*j*_ due to its interaction with the *transmitter p_i_*. The impact of *p*_*i*_ on *p*_*j*_ increases with the magnitude of *τ*_*i*→*j*_.
5. The *τ*_*i*→*j*_ is a relative measure which quantifies the information transfer from *p*_*i*_ to *p*_*j*_ with respect to the endogenous and noise-induced changes of the latter. However, it cannot be used to compare the information flow rate from *p*_*i*_ to *p*_*j*_ with that from *p*_*j*_ to *p*_*i*_. This is due to the fact that the normalization of *τ*_*i*→*j*_ depends on the entropy changes of *p*_*j*_, while the normalization of *τ*_*j*→*i*_ depends on the entropy changes of *p*_*i*_. The comparison of the reverse information flows between *p*_*i*_ and *p*_*j*_ should thus be based on the non-normalized coefficients *T*_*i*→*j*_ and *T*_*j*→*i*_(*Liang, 2015*).
6. The information flow rates (normalized and non-normalized) can be calculated without requiring (i) the estimation of conditional probability distributions (ii) stationarity assumptions (iii) Gaussian distribution of the fluctuations or (iv) a specific model structure.

### Non-parametric testing of normalized information flow rate

To calculate *τ*_*i*→*j*_ for each individual *l* = 1, … *L*, we use all the time points in the series 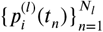, for the source locations *i* = 1, …, 15. Each series represents the strength of the current dipole moment at location *i*. All the time series for the same individual (indexed by *l*) have the same length *N*_*l*_ which varies between 67845 and 114304 points.

In order to infer connectivity patterns, it is necessary to know if the estimated values *τ*_*i*→*j*_ are statistically significant. Each estimate of an inter-dipole *τ*_*i*→*j*_ is a statistic, i.e., a random variable that fluctuates between samples. If the sampling distribution of the statistic is known, the significance of a particular estimate can be assessed using a suitably constructed parametric statistical test. In the case of *T*_*i*→*j*_ such a test can be constructed (*Liang, 2014*). For *τ*_*i*→*j*_, however, the sampling distribution is not known. In this case, it is possible to apply *non-parametric permutation testing* in the spirit used by Lachaux *et al.* to quantify the significance of phase locking values (*Lachaux et al., 1999*; *Bastos et al., 2015*). The goal of non-parametric permutation testing is to determine the probability that the observed test statistic could have been realized if the null hypothesis (i.e., zero information flow) were true. This, in turn, allows us to conclude if an estimated information flow rate is statistically significant: a very small probability (*p*-value) implies that the observed deviation is not likely under the null hypothesis (*Maris and Oostenveld, 2007*; *Cohen, 2014*).

The test statistic that we use is the normalized information flow rate from series *p*_*i*_ to series *p*_*j*_, for *i ≠ j* = 1, …, 15. The null hypothesis that we test is that there is no information flow between series *p*_*i*_ and *p*_*j*_. We generate *M*_*s*_ = 1000 randomized states 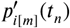, where *m* = 1, …, *M_s_* and *n* = 1, …, *N*, from *each* transmitter time series *p*_*i*_. Each randomized state is obtained by scrambling (by means of random permutations) the *N* time points of *p*_*i*_. The permutation destroys the temporal ordering of *p*_*i*_ and consequently any patterns of information flow from 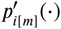 to 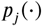. Hence, the estimated *τ*_*i*[*m*]→*j*_ values based on the shuffled time series *p*_*i*_ do not represent meaningful information flow.

The *p*-value of the statistic *τ*_*i*→*j*_ is defined as the percentage of times — calculated over *M*_*s*_ permutation states — that the randomized information flow rate *τ*_*i*[*m*]→*j*_ is more extreme than *τ*_*i*→*j*_ (i.e., larger than *τ*_*i*→*j*_ if *τ*_*i*→*j*_ > 0 and smaller than *τ*_*i*→*j*_ if *τ*_*i*→*j*_ < 0). A high *p*-value would indicate that the null hypothesis cannot be rejected. In contrast, a low *p*-value would provide support for the alternative hypothesis (i.e., that there is significant information flow from *p*_*i*_ to *p*_*j*_). The observed value *τ*_*i*→*j*_ is then considered as statistically significant, if the respective *p*-value is below a specified significance level (typically 0.1%–5%).

Based on all the simulations performed for the entire cohort of 32 individuals, we find that the magnitude of the information flow rates between the randomly permuted *transmitter* current dipoles and the *receiver* current dipoles are all contained in the interval [−5.5, 5.0] × 10^−4^. Turning to the inter-dipole information flow rates calculated from the EEG data, we find that all except for 11 out of the 210×32=6720 dipole pairs show information flow rates outside the above interval. In fact, the majority of the normalized information flow rates are two orders or more higher in magnitude. Hence, given the size of the above confidence interval, we can conclude that most of the observed *τ*_*i*→*j*_ are statistically significant even at the *p* = 0.1% level.

The above result indicates a low-level global connectivity linking most of the brain regions in the resting state. However, small normalized information flow rates, albeit statistically significant, imply that the contribution of the respective entropy flow rate (information flow) from the *transmitter* dipole to the *receiver* dipole is very small compared to the intrinsic entropy changes in the *receiver* dipole. This argument motivates the introduction of an arbitrary threshold that can be used to count the more important connections.

### Impact of differencing scheme on connectivity

As stated following the definition of inter-dipole information flow rate, the estimation of the first-order derivatives is based on finite differences (cf. *Equation 4*). The finite differencing, as shown in *Equation 4*, can be accomplished by means of different time steps equal to *kΔt*. Typically, *k* = 1 or *k* = 2 is used (*Liang, 2014*). We have conducted our analysis with *k* = 2, since this choice tends to reduce the impact of occasional large spikes (e.g., jumps) in the EEG time series on the information flow rate. Using a larger value of *k* results in effective smoothing of the EEG time series which encroaches on the upper end of the frequency band that is generally of interest in resting state studies. Hence, we did not consider values of *k* higher than two.

We have experimented with synthetic data obtained from the simulation of two coupled stochastic differential equations for which *T*_*i*→*j*_ admits explicit expressions (*Liang, 2014*). We used similar length *N* = 60000 − 80000) for the synthetic time series as that of the EEG series and a number of repetitions equal to the number of individuals in the study (*L* = 32). Our results show practically no difference between the mean *T*_*i*→*j*_ estimated from the time series whether *k* = 1 or *k* = 2 is used.

In the case of the source-reconstructed EEG data, we repeated the entire analysis using *k* = 1. This leads to fewer active connections, i.e., 1904 instead of 2821 for *k* = 2 shown in *Figure 8*. On the other hand, the correlation coefficient between the spatial distribution of the frequency of active connections *n*_*i*→*j*_(0.05) for *k* = 1 and for *k* = 2 is equal to 0.89. This indicates that the distribution of active connections is highly correlated between *k* = 1 and *k* = 2. Based on our arbitrary threshold *τ*_*c*_ = 0.05, we determine 42 active connections 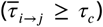 for the *k* = 1 scheme versus 92 active connections for the *k* = 2 scheme. In spite of the fact that fewer active connections appear for *k* = 1, the overall pattern of information flow, as delineated by the thirty connections with the highest 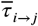, remains unchanged.

## Acknowledgments

DTH acknowledges useful electronic correspondence with X. San Liang regarding the definition and interpretation of the information flow rate coefficient.

## Appendix 1

Herein we show how the two entropic components involved, in addition to |*T*_*i*→*j*_|, in the normalization term *Z*_*i*→*j*_, can be estimated from the data. Analysis based on the theory of dynamical systems leads to the following expressions for the rates of change of the entropic components 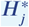 and 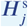 (*Liang, 2008*)

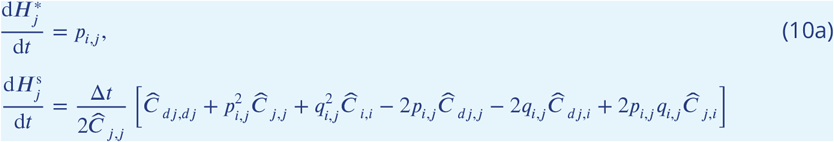

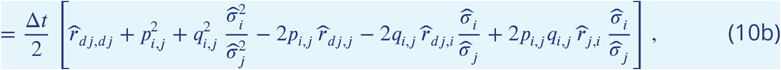

where the entropy transfer elements *p_i,j_*, *q*_*i,j*_ are given by the following functions of the inter-dipole covariance coefficients

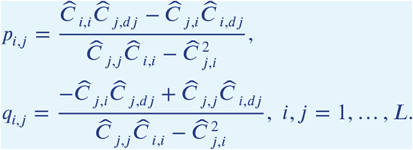

Using the definition in *Equation 2* for the correlation coefficient and the definition in *Equation 3* for the cross-correlation coefficient, the elements *p_i,j_* and *q*_*i,j*_ can be expressed using correlation coefficients *r* instead of inter-dipole covariances *C* as follows (for *i, j* = 1, …, *N_s_*):

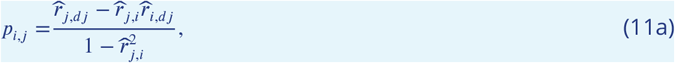

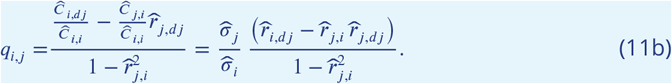

## Footnotes

1 http://www.besa.de

## References

Agcaoglu O, Miller R, Mayer AR, Hugdahl K, Calhoun VD. Lateralization of resting state networks and relationship to age and gender. NeuroImage. 2015; 104:310–325. doi: 10.1016/j.neuroimage.2014.09.001.

Baccalá LA, Sameshima K. Partial directed coherence: a new concept in neural structure determination. Biological cybernetics. 2001; 84(6):463–474. doi: 10.1007/PL00007990.

Barnett L, Barrett AB, Seth AK. Granger causality and transfer entropy are equivalent for Gaussian variables. Physical Review Letters. 2009 Dec; 103:238701. doi: 10.1103/PhysRevLett.103.238701.

Bastos AM, Schoffelen JM. A tutorial review of functional connectivity analysis methods and their interpretational pitfalls. Frontiers in Systems Neuroscience. 2015; 9. doi: 10.3389/fnsys.2015.00175.

Bastos AM, Vezoli J, Conrado AB, Schoffelen JM, Oostenveld R, Dowdall JR, De Weerd P, Kennedy H, Pascal F. Visual areas exert feedforward and feedback influences through distinct frequency channels. Neuron. 2015; 85(2):390–401. doi: 10.1016/j.neuron.2014.12.018.

BESA Wiki, BESA Research Montage Editor; 2018. http://wiki.besa.de/index.php?title=BESA_Research_Montage_Editor#Virtual_Standard_Montages, [Online; accessed 31-May-2018].

Blanco S, Garcia H, Quiroga RQ, Romanelli L, Rosso OA. Stationarity of the EEG series. IEEE Engineering in Medicine and Biology Magazine. 1995; 14(4):395–399. doi: 10.1109/51.395321.

Borich M, Babul AN, Yuan PH, Boyd L, Virji-Babul N. Alterations in resting-state brain networks in concussed adolescent athletes. Journal of Neurotrauma. 2015; 32(4):265–271. doi: 10.1089/neu.2013.3269.

Bossomaier T, Barnett L, Harré M, Lizier JT. An Introduction to Transfer Entropy: Information Flow in Complex Systems. Cham, Switzerland: Springer; 2016.

Bressler SL, Seth AK. Wiener–Granger causality: A well established methodology. NeuroImage. 2011; 58(2):323–329. doi: 10.1016/j.neuroimage.2010.02.059.

Chen Y, Rangarajan G, Feng J, Ding M. Analyzing multiple nonlinear time series with extended Granger causality. Physics Letters A. 2004; 324(1):26–35. doi: 10.1016/j.physleta.2004.02.032.

Cohen MX. Analyzing Neural Time Series Data: Theory and Practice. MIT Press; 2014.

Ding M, Chen Y, Bressler SL. Granger Causality: Basic Theory and Application to Neuroscience. In: Handbook of Time Series Analysis John Wiley & Sons, Ltd; 2006.p. 437–460. doi: 10.1002/9783527609970.ch17.

Emberson LL, Richards JE, Aslin RN. Top-down modulation in the infant brain: Learning-induced expectations rapidly affect the sensory cortex at 6 months. Proceedings of the National Academy of Sciences. 2015; 112(31):9585–9590. doi: 10.1073/pnas.1510343112.

Fair DA, Cohen AL, Dosenbach NUF, Church JA, Miezin FM, Barch DM, Raichle ME, Petersen SE, Schlaggar BL. The maturing architecture of the brain’s default network. Proceedings of the National Academy of Sciences. 2008; 105(10):4028–4032. doi: 10.1073/pnas.0800376105.

Friston KJ. Functional and effective connectivity: a review. Brain Connectivity. 2011; 1(1):13–36. doi: 10.1089/brain.2011.0008.

Friston KJ. Functional and effective connectivity in neuroimaging: a synthesis. Human brain mapping. 1994; 2(1-2):56–78. doi: 10.1002/hbm.460020107.

Friston KJ, Harrison L, Penny W. Dynamic causal modelling. NeuroImage. 2003; 19(4):1273–1302. doi: 10.1016/S1053-8119(03)00202-7.

Gogtay N, Giedd JN, Lusk L, Hayashi KM, Greenstein D, Vaituzis AC, Nugent TF, Herman DH, Clasen LS, Toga AW, Rapoport JL, Thompson PM. Dynamic mapping of human cortical development during childhood through early adulthood. Proceedings of the National Academy of Sciences. 2004; 101(21):8174–8179. doi: 10.1073/pnas.0402680101.

Greicius MD, Supekar K, Menon V, Dougherty RF. Resting-state functional connectivity reflects structural connectivity in the default mode network. Cerebral Cortex. 2009; 19(1):72–78. doi: 10.1093/cercor/bhn059.

Gutzwiller MC. Chaos in Classical and Quantum Mechanics, vol. 1 of Interdisciplinary Applied Mathematics. New York: Springer; 1990.

Hesse W, Möller E, Arnold M, Schack B. The use of time-variant EEG Granger causality for inspecting directed interdependencies of neural assemblies. Journal of Neuroscience Methods. 2003; 124(1):27–44. doi: 10.1016/S0165-0270(02)00366-7.

Hillebrand A, Tewarie P, van Dellen E, Yu M, Carbo EWS, Douw L, Gouw AA, van Straaten ECW, Stam CJ. Direction of information flow in large-scale resting-state networks is frequency-dependent. Proceedings of the National Academy of Sciences. 2016; 113(14):3867–3872. doi: 10.1073/pnas.1515657113.

Hirayama J, Hyvärinen A, Kiviniemi V, Kawanabe M, Yamashita O. Characterizing variability of modular brain connectivity with constrained principal component analysis. PLoS ONE. 2016 12; 11(12):1–24. doi: 10.1371/journal.pone.0168180.

Horwitz B. The elusive concept of brain connectivity. NeuroImage. 2003; 19(2):466–470. doi: 10.1016/S1053-8119(03)00112-5.

Hutchison RM, Womelsdorf T, Allen EA, Bandettini PA, Calhoun VD, Corbetta M, Della Penna S, Duyn JH, Glover GH, Gonzalez-Castillo J, et al. Dynamic functional connectivity: promise, issues, and interpretations. NeuroImage. 2013; 80:360–378. doi: 10.1016/j.neuroimage.2013.05.079.

Kamiński M, Ding M, Truccolo WA, Bressler SL. Evaluating causal relations in neural systems: Granger causality, directed transfer function and statistical assessment of significance. Biological Cybernetics. 2001; 85(2):145–157. doi: 10.1007/s004220000235.

Kaplan AY, Fingelkurts AA, Fingelkurts AA, Borisov SV, Darkhovsky BS. Nonstationary nature of the brain activity as revealed by EEG/MEG: Methodological, practical and conceptual challenges. Signal Processing. 2005; 85(11):2190–2212. doi: 10.1016/j.sigpro.2005.07.010.

Klonowski W. Everything you wanted to ask about EEG but were afraid to get the right answer. Nonlinear Biomedical Physics. 2009; 3(1):2. doi: 10.1186/1753-4631-3-2.

Lachaux JP, Rodriguez E, Martinerie J, Varela FJ. Measuring phase synchrony in brain signals. Human Brain Mapping. 1999; 8(4):194–208. doi: 10.1002/(SICI)1097-0193(1999)8:4\%3C194::AID-HBM4\%3E3.0.CO;2-C.

Lebel C, Beaulieu C. Longitudinal development of human brain wiring continues from childhood into adulthood. Journal of Neuroscience. 2011; 31(30):10937–10947. doi: 10.1523/JNEUROSCI.5302-10.2011.

Li J, Wang ZJ. Controlling the false discovery rate of the association/causality structure learned with the PC algorithm. Journal of Machine Learning Research. 2009 Jun; 10:475–514.

Liang XS. Information flow within stochastic dynamical systems. Physical Review E. 2008; 78(3):031113. doi: 10.1103/PhysRevE.78.031113.

Liang XS. The Liang-Kleeman information flow: Theory and applications. Entropy. 2013; 15(1):327–360. doi: 10.3390/e15010327.

Liang XS. Local predictability and information flow in complex dynamical systems. Physica D: Nonlinear Phenomena. 2013; 248:1–15. doi: 10.1016/j.physd.2012.12.011.

Liang XS. Unraveling the cause-effect relation between time series. Physical Review E. 2014; 90(5):052150. doi: 10.1103/PhysRevE.90.052150.

Liang XS. Normalizing the causality between time series. Physical Review E. 2015; 92(2):022126. doi: 10.1103/PhysRevE.92.022126.

Liang XS. Information flow and causality as rigorous notions ab initio. Physical Review E. 2016; 94(5):052201. doi: 10.1103/PhysRevE.94.052201.

Liang XS. Causation and information flow with respect to relative entropy. Chaos: An Interdisciplinary Journal of Nonlinear Science. 2018; 28(7):075311. doi: 10.1063/1.5010253.

Liang XS, Kleeman R. Information Transfer between Dynamical System Components. Physica Review Letters. 2005; 95:244101. doi: 10.1103/PhysRevLett.95.244101.

Lin P, Yang Y, Gao J, De Pisapia N, Ge S, Wang X, Zuo CS, Levitt JJ, Niu C. Dynamic default mode network across different brain states. Scientific Reports. 2017; 7:46088. doi: 10.1038/srep46088.

Lindquist MA. The statistical analysis of fMRI data. Statistical Science. 2008; 23(4):439–464. doi: 10.1214/09-STS282.

Liu Q, Farahibozorg S, Porcaro C, Wenderoth N, Mantini D. Detecting large-scale networks in the human brain using high-density electroencephalography. Human Brain Mapping. 2017; 38(9):4631–4643. doi: 10.1002/hbm.23688.

Liu Y, Aviyente S. Quantification of effective connectivity in the brain using a measure of directed information. Computational and Mathematical Methods in Medicine. 2012; 2012. doi: 10.1155/2012/635103.

Marinazzo D, Liao W, Chen H, Stramaglia S. Nonlinear connectivity by Granger causality. NeuroImage. 2011; 58(2):330–338. doi: 10.1016/j.neuroimage.2010.01.099.

Maris E, Oostenveld R. Nonparametric statistical testing of EEG- and MEG-data. Journal of Neuroscience Methods. 2007; 164(1):177–190. doi: 10.1016/j.jneumeth.2007.03.024.

Marshall WJ, Lackner CL, Marriott P, Santesso DL, Segalowitz SJ. Using phase shift Granger causality to measure directed connectivity in EEG recordings. Brain Connectivity. 2014; 4(10):826–841. doi: 10.1089/brain.2014.0241.

McLntosh AR, Gonzalez-Lima F. Structural equation modeling and its application to network analysis in functional brain imaging. Human Brain Mapping. 1994; 2(1-2):2–22. doi: 10.1002/hbm.460020104.

Michels L, Muthuraman M, Lüchinger R, Martin E, Anwar AR, Raethjen J, Brandeis D, Siniatchkin M. Developmental changes of functional and directed resting-state connectivities associated with neuronal oscillations in EEG. NeuroImage. 2013; 81:231–242. doi: 10.1016/j.neuroimage.2013.04.030.

Mill RD, Bagic A, Bostan A, Schneider W, Cole MW. Empirical validation of directed functional connectivity. NeuroImage. 2017; 146:275–287. doi: 10.1016/j.neuroimage.2016.11.037.

Miskovic V, Ma X, Chou CA, Fan M, Owens M, Sayama H, Gibb BE. Developmental changes in spontaneous electrocortical activity and network organization from early to late childhood. NeuroImage. 2015; 118:237–247. doi: 10.1016/j.neuroimage.2015.06.013.

Muetzel RL, Blanken LME, Thijssen S, van der Lugt A, Jaddoe VWV, Verhulst FC, Tiemeier H, White T. Resting-state networks in 6-to-10 year old children. Human Brain Mapping. 2016; 37(12):4286–4300. doi: 10.1002/hbm.23309.

Muthuraman M, Moliadze V, Mideksa KG, Anwar AR, Stephani U, Deuschl G, Freitag CM, Siniatchkin M. EEG-MEG integration enhances the characterization of functional and effective connectivity in the resting state network. PLoS ONE. 2015; 10(10):e0140832. doi: 10.1371/journal.pone.0140832.

Paus T. Mapping brain maturation and cognitive development during adolescence. Trends in Cognitive Sciences. 2005; 9(2):60–68. doi: 10.1016/j.tics.2004.12.008.

Perrin F, Pernier J, Bertrand O, Echallier JF. Spherical splines for scalp potential and current density mapping. Electroencephalography and Clinical Neurophysiology. 1989; 72(2):184–187. doi: 10.1016/0013-4694(89)90180-6.

Porter S, Torres IJ, Panenka W, Rajwani Z, Fawcett D, Hyder A, Virji-Babul N. Changes in brain-behavior relationships following a 3-month pilot cognitive intervention program for adults with traumatic brain injury. Heliyon. 2017; 3(8):e00373. doi: 10.1016/j.heliyon.2017.e00373.

Raichle ME, MacLeod AM, Snyder AZ, Powers WJ, Gusnard DA, Shulman GL. A default mode of brain function. Proceedings of the National Academy of Sciences. 2001; 98(2):676–682. doi: 10.1073/pnas.98.2.676.

Raichle ME, Mintun MA. Brain work and brain imaging. Annual Reviews of Neuroscience. 2006; 29:449–476. doi: 10.1146/annurev.neuro.29.051605.112819.

Roebroeck A, Formisano E, Goebel R. Mapping directed influence over the brain using Granger causality and fMRI. NeuroImage. 2005; 25(1):230–242. doi: https://doi.org/10.1016/j.neuroimage.2004.11.017.

Rotem-Kohavi N, Hilderman CGE, Liu A, Makan N, Wang JZ, Virji-Babul N. Network analysis of perception-action coupling in infants. Frontiers in Human Neuroscience. 2014; 8:209. doi: 10.3389/fnhum.2014.00209.

Rotem-Kohavi N, Oberlander TF, Virji-Babul N. Infants and adults have similar regional functional brain organization for the perception of emotions. Neuroscience Letters. 2017; 650(Supplement C):118–125. doi: 10.1016/j.neulet.2017.04.031.

Rubinov M, Sporns O. Complex network measures of brain connectivity: Uses and interpretations. NeuroImage. 2010; 52(3):1059–1069. doi: 10.1016/j.neuroimage.2009.10.003.

Sakkalis V. Review of advanced techniques for the estimation of brain connectivity measured with EEG/MEG. Computers in Biology and Medicine. 2011; 41(12):1110–1117. doi: 10.1016/j.compbiomed.2011.06.020.

Salvador R, Anguera M, Gomar J, Bullmore E, Pomarol-Clotet E. Conditional mutual information maps as descriptors of net connectivity levels in the brain. Frontiers in Neuroinformatics. 2010; 4:115–123. doi: 10.3389/fninf.2010.00115.

Scherg M, Ille N, Bornfleth H, Berg P. Advanced tools for digital EEG review: Virtual source montages, whole-head mapping, correlation, and phase analysis. Journal of Clinical Neurophysiology. 2002; 19(2):91–112.

Schreiber T. Measuring information transfer. Physical Review Letters. 2000; 85(2):461–464. doi: 10.1103/Phys-RevLett.85.461.

Seth A. Granger causality. Scholarpedia. 2007; 2(7):1667. revision #91329.

Seth AK, Barrett AB, Barnett L. Granger causality analysis in neuroscience and neuroimaging. Journal of Neuroscience. 2015; 35(8):3293–3297. doi: 10.1523/JNEUROSCI.4399-14.2015.

Shaw P, Kabani NJ, Lerch JP, Eckstrand K, Lenroot R, Gogtay N, Greenstein D, Clasen L, Evans A, Rapoport JL, Giedd JN, Wise SP. Neurodevelopmental trajectories of the human cerebral cortex. Journal of Neuroscience. 2008; 28(14):3586–3594. doi: 10.1523/JNEUROSCI.5309-07.2008.

Shovon MHI, Nandagopal DN, Vijayalakshmi R, Du JT, Cocks B. Transfer entropy and information flow patterns in functional brain networks during cognitive activity. In: International Conference on Neural Information Processing Springer; 2014. p. 1–10.

Smit DJA, Boersma M, Schnack HG, Micheloyannis S, Boomsma DI, Pol HEH, Stam CJ, de Geus EJC. The brain matures with stronger functional connectivity and decreased randomness of its network. PLoS ONE. 2012; 7(5):e36896. doi: 10.1371/journal.pone.0036896.

Snyder AZ, Raichle ME. A brief history of the resting state: the Washington University perspective. NeuroImage. 2012; 62(2):902–910. doi: 10.1016/j.neuroimage.2012.01.044.

Sporns O. Networks of the Brain. MIT Press; 2011.

Stam CJ, Van Straaten ECW. The organization of physiological brain networks. Clinical Neurophysiology. 2012; 123(6):1067–1087. doi: 10.1016/j.clinph.2012.01.011.

Taylor MJ, Khan SC. Top-down modulation of early selective attention processes in children. International Journal of Psychophysiology. 2000; 37(2):135–147. doi: 10.1016/S0167-8760(00)00084-2.

Toga AW, Thompson PM, Mori S, Amunts K, Zilles K. Towards multimodal atlases of the human brain. Nature Reviews Neuroscience. 2006; 7(12):952–966. doi: 10.1038/nrn2012.

Vicente R, Wibral M, Lindner M, Pipa G. Transfer entropy—a model-free measure of effective connectivity for the neurosciences. Journal of Computational Neuroscience. 2011; 30(1):45–67. doi: 10.1007/s10827-010-0262-3.

Van de Ville D, Britz J, Michel CM. EEG microstate sequences in healthy humans at rest reveal scale-free dynamics. Proceedings of the National Academy of Sciences. 2010; 107(42):18179–18184. doi: 10.1073/pnas.1007841107.

Virji-Babul N, Hilderman CGE, Makan N, Liu A, Smith-Forrester J, Franks C, Wang ZJ. Changes in functional brain networks following sports-related concussion in adolescents. Journal of Neurotrauma. 2014; 31(23):1914–1919. doi: /10.1089/neu.2014.3450.

Yuan H, Ding L, Zhu M, Zotev V, Phillips R, Bodurka J. Reconstructing large-scale brain resting-state networks from high-resolution EEG: spatial and temporal comparisons with fMRI. Brain Connectivity. 2016; 6(2):122–135. doi: 10.1089/brain.2014.0336.

